# CXCR2 inhibition enables NASH-HCC immunotherapy

**DOI:** 10.1101/2022.02.24.481779

**Authors:** Jack Leslie, John B. G. Mackey, Thomas Jamieson, Erik Ramon-Gil, Thomas M. Drake, Frédéric Fercoq, William Clark, Kathryn Gilroy, Ann Hedley, Colin Nixon, Saimir Luli, Maja Laszczewska, Roser Pinyol, Roger Esteban-Fabró, Catherine E. Willoughby, Philipp K. Haber, Carmen Andreu-Oller, Mohammad Rahbari, Chaofan Fan, Dominik Pfister, Shreya Raman, Niall Wilson, Miryam Müller, Amy Collins, Daniel Geh, Andrew Fuller, David McDonald, Gillian Hulme, Andrew Filby, Xabier Cortes-Lavaud, Noha-Ehssan Mohamed, Catriona A. Ford, Ximena L. Raffo Iraolagoitia, Amanda J. McFarlane, Misiti McCain, Rachel Ridgway, Edward W. Roberts, Simon T. Barry, Gerard J. Graham, Mathias Heikenwalder, Helen L. Reeves, Josep M. Llovet, Leo M. Carlin, Thomas G. Bird, Owen J. Sansom, Derek A. Mann

## Abstract

**Objective:** Hepatocellular carcinoma (HCC) is increasingly associated with non-alcoholic steatohepatitis (NASH). HCC immunotherapy offers great promise; however, recent data suggests NASH-HCC may be less sensitive to conventional immune checkpoint inhibition (ICI). We hypothesised that targeting neutrophils using a CXCR2 small molecule inhibitor may sensitise NASH-HCC to ICI therapy.

**Design:** Neutrophil infiltration was characterised in human HCC and mouse models of HCC. Late-stage intervention with anti-PD1 and/or a CXCR2 inhibitor was performed in murine models of NASH-HCC. The tumour immune microenvironment was characterised by imaging mass cytometry, RNA-seq and flow cytometry.

**Results:** Neutrophils expressing CXCR2, a receptor crucial to neutrophil recruitment in acute-injury, are highly represented in human NASH-HCC. In models of NASH-HCC lacking response to ICI, the combination of a CXCR2 antagonist with anti-PD1 suppressed tumour burden and extended survival. Combination therapy increased intratumoral XCR1^+^ dendritic cell activation and CD8^+^ T cell numbers which are associated with anti-tumoral immunity, this was confirmed by loss of therapeutic effect upon genetic impairment of myeloid cell recruitment, neutralisation of the XCR1-ligand XCL1 or depletion of CD8^+^ T cells. Therapeutic benefit was accompanied by an unexpected increase in tumour-associated neutrophils (TANs) which switched from a pro-tumor to anti-tumour progenitor-like neutrophil phenotype. Reprogrammed TANs were found in direct contact with CD8^+^ T cells in clusters that were enriched for the cytotoxic anti-tumoural protease granzyme B. Neutrophil reprogramming was not observed in the circulation indicative of the combination therapy selectively influencing TANs.

**Conclusion:** CXCR2-inhibition induces reprogramming of the tumour immune microenvironment that promotes ICI in NASH-HCC.

**Significance of this study:** **What is already known on this subject?**

- Immune checkpoint inhibition therapy is emerging as a promising new therapy for the treatment of advanced hepatocellular carcinoma (HCC).
- Only a minority of HCC patients will respond to immune checkpoint inhibition (ICI) therapy and recent data suggest that HCC on the background of NASH may have reduced sensitivity to this treatment strategy.
- Neutrophils are a typical myeloid component of the liver in NASH and are found either within the HCC tumour microenvironment or in a peritumoural location.
- Neutrophils have considerable phenotypic plasticity and can exist in both tumour promoting and tumour suppressing states.
- Neutrophils may have the ability to influence ICI therapy.

**What are the new findings?**

- CXCR2^+^ neutrophils are found in human NASH and within the tumour of both human and mouse models of NASH-HCC.
- The resistance of NASH-HCC to anti-PD1 therapy is overcome by co-treatment with a CXCR2 small molecule inhibitor, with evidence of reduced tumour burden and extended survival.
- Anti-PD1 and CXCR2 inhibitor combine to selectively reprogramme tumour-associated neutrophils (TANs) from a pro- to an anti-tumour phenotype.
- Reprogrammed TANs proliferate locally within Granzyme B^+^ immune clusters that contain physically associating CD8^+^ T cells and antigen presenting cells.
- Conventional XCR1^+^ dendritic cells (cDC1s) are found to be elevated in anti-PD1 and CXCR2 inhibitor treated HCCs and together with CD8^+^ T cells are required for therapeutic benefit.

**How might it impact on clinical practice in the foreseeable future?**

- TANs can be selectively manipulated to adopt an anti-tumour phenotype which unlocks their potential for cancer therapy. The ability of CXCR2 antagonism to combine with ICI therapy to bring about enhanced therapeutic benefit in NASH-HCC (and potentially in HCC of other aetiologies) warrents clinical investigation.

## Introduction

Primary liver cancer is emerging globally as one of the most common and deadly malignancies with 905,000 new diagnosed cases and 830,000 deaths recorded in 2020^1^. Hepatocellular carcinoma (HCC) accounts for up to 85% of primary liver cancers and develops on the background of chronic liver disease (CLD) caused by persistent virological (HBV and HCV) or non-virological liver damage. Due to the increasing prevalence of obesity and the metabolic syndrome a high proportion of HCC is now attributed to non-alcoholic steatohepatitis (NASH), identified as the most common risk factor for HCC in UK and USA^2,3^.

Possible curative options for HCC such as tumour resection, liver transplant or ablation are at present limited to a minority of patients who are diagnosed at an early stage of the disease^4^. For more advanced HCC, approved systemic therapies include multikinase inhibitors and anti-VEGF agents. More recently, immune checkpoint inhibition (ICI) has emerged as a therapeutic modality in HCC with PD1 antibodies (nivolumab and pembrolizumab) being approved, and a combination of anti-PDL1 (atezolizumab) with anti-VEGF (bevacizumab) now being first-line treatment for advanced HCC^5–7^. However, only a minority (up to 30%) of HCC patients respond to immunotherapy^5–8^. Moreover, it was recently reported that HCC on the background of NASH is less responsive to immunotherapy due to a NASH-induced alteration in the immune components of the liver and in particular an expansion in numbers of exhausted CD8^+^PD1^+^ T cells, that appear to promote, rather than suppress, NASH-HCC^9,10^. Therefore, advanced therapeutic strategies for HCC will require a deeper appreciation of the complex immune landscape of the tumour microenvironment, and in particular, should also take into account the influence that the background liver pathology may have on the numbers, regional distributions, phenotypes and activities of key immune cell types of relevance to cancer growth.

Recent use of imaging mass cytometry and single cell sequencing to probe the cellular constituents of human HCC revealed considerable heterogeneity within the tumour microenvironment with intratumoural region-specific distributions of immune cells^11^. Regions with evidence of less aggressive cancer and ongoing liver damage (fibrogenesisis) were enriched for CD8^+^ T cells, B cells and CD11b^+^CD15^+^GranzymeB^+^ neutrophils. When considering the growing evidence for both pro- and anti-tumour functions for neutrophils in a variety of cancers^12,13^ including HCC^14^ we were interested to determine if modulation of neutrophil biology within the tumour microenvironment would influence the resistance of NASH-HCC to anti-PD1 immunotherapy.

Here, we determined that the CXC chemokine receptor CXCR2 is almost exclusively located on neutrophils in human and mouse NASH-HCC. This finding led us to ask if antagonism of CXCR2 can combine with anti-PD1 to overcome resistance of NASH-HCC to immunotherapy. Our findings suggest that this combination therapy reprogrammes the phenotype of tumour neutrophils and enhances their association with CD8^+^ T cells and conventional dendritic cells (cDC). Reshaping the tumour immune microenvironment was associated with a T cell- and DC-dependent reduction in tumour burden and increased survival. We propose that combined neutrophil phenotype modification and ICI may achieve improved outcomes in NASH-HCC.

### Pro-tumour CXCR2^+^ neutrophils associate with NASH-HCC resistance to anti-PD1 immunotherapy

To investigate the immunological determinants of unresponsiveness of NASH-HCC to anti-PD1 therapy we designed an orthotopic mouse model using the Hep-53.4 HCC cell line, which was selected due to its high mutational burden (**Supplementary Data Fig. 1a-c**). On the background of steatosis induced by a modified diet of high sugar and fat, we observed weight gain and larger tumours compared to non-steatotic controls (**Supplementary Data Fig. 1d-g**). Tumours in non-steatotic controls were responsive to anti-PD1 therapy, however, anti-PD1 showed no benefit on tumour burden, survival, steatosis, proliferation or immune cell infiltration in steatotic mice (**Fig. 1a-f** and **Supplementary Data Fig. 1h-k**). For an additional autochthonous model, we employed either Diethylnitrosamine (DEN) alone or in combination with the American lifestyle induced obesity syndrome diet (DEN/ALIOS), the latter to establish HCC on a background of NASH^15,16^ (**Supplementary Data Fig. 1l-o**). Anti-PD1 responsiveness was observed in DEN mice fed a control diet, whereas anti-PD1 therapy had no effect on tumour burden, proliferation, or steatosis and animal survival when mice were fed the ALIOS diet (**Fig. 1g-l** and **Supplementary Data Fig. 1p, q**). However, F4/80^+^ and CD3^+^ immune cell infiltrates were increased in anti-PD1 treated ALIOS fed mice (**Supplementary Data Fig. 1r, s**) indicative of anticipated alterations in tumour immunity.

**Figure 1.**
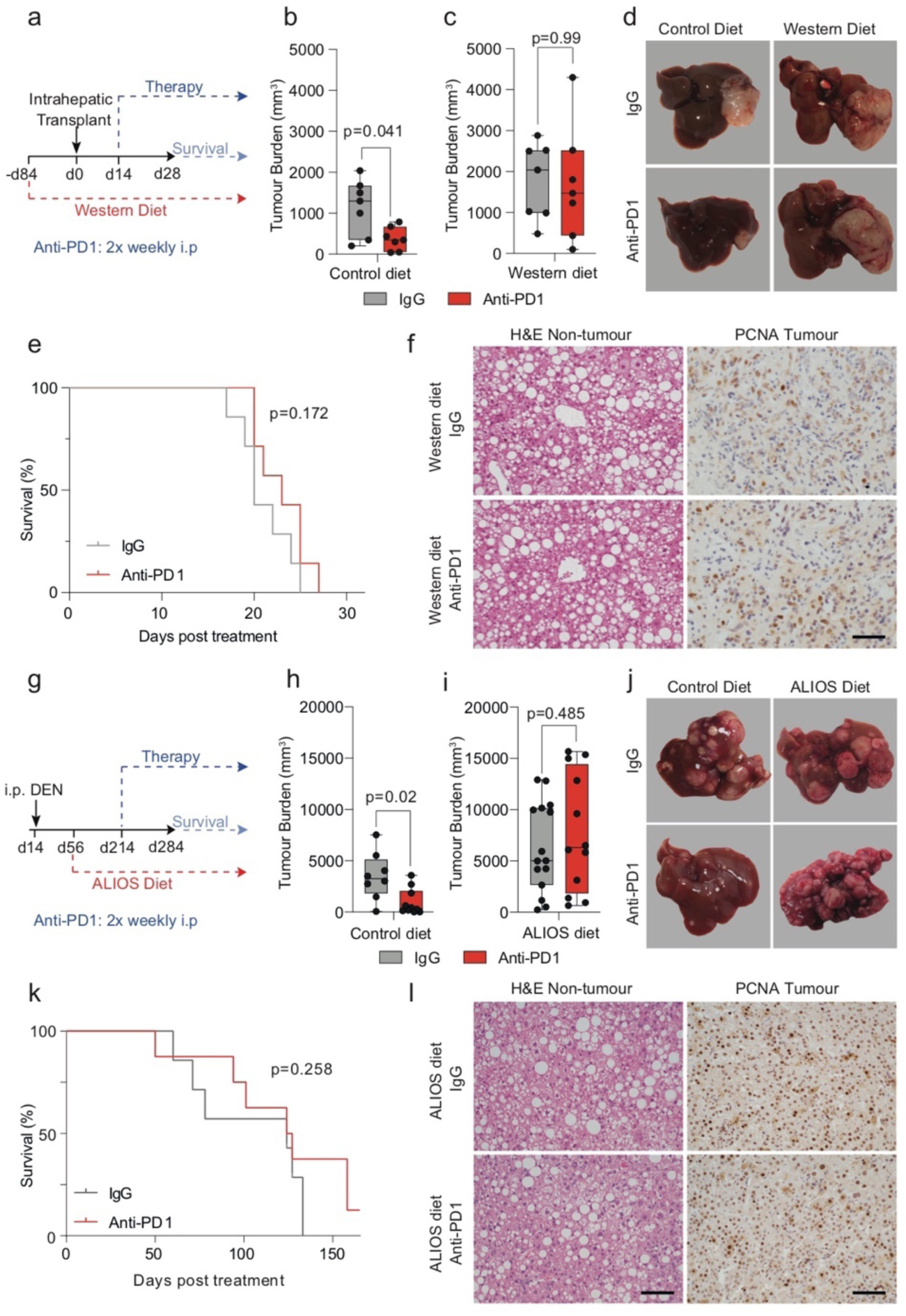
NASH-HCC is resistant to anti-PD1 immunotherapy. a) Timeline schematic of the NASH-HCC model based on intra-hepatic injection of Hep53.4 HCC cells into C57BL/6 mice fed a control or Western diet. Cells were implaned 84 days after diet initiation, and anti-PD1 therapy was administered from day 14 post-transplantation. b) Quantification of tumour burden (mm^3^) at day 28 post-intrahepatic injection for orthotopic HCC mice fed a control diet and treated with IgG-control or anti-PD1. n=7 mice per condition. Mann-Whitney *U*-test. c) Quantification of tumour burden (mm^3^) at day 28 post-intrahepatic injection for orthotopic NASH-HCC mice fed a Western diet and treated with IgG-control or anti-PD1. n=7 mice per condition. Mann-Whitney *U*-test. d) Representative images of livers at day 28 post-intrahepatic injection for orthotopic NASH-HCC mice fed a control or Western diet and treated with IgG-control or anti-PD1 from day 14. n=7 mice per condition. e) Survival plot in orthotopic NASH-HCC mice fed a Western diet and treated with IgG-control or anti-PD1. n=7 mice per condition. Log-rank (Mantel-Cox) test. f) Representative images of H&E-stained non-tumour livers and PCNA-stained tumours from orthotopic NASH-HCC mice fed a Western diet and treated with IgG-control or anti-PD1 at day 28 post-intrahepatic injection. n=7 mice per condition. Scale bar = 100 µm. g) Timeline schematic for the administration of Diethylnitrosamine (DEN) and the American lifestyle induced obesity syndrome (ALIOS) diet used to generate the autochthonous DEN/ALIOS NASH-HCC model with therapy timelines indicated. h) Quantification of tumour burden (mm^3^) at day 284 for DEN mice fed a control diet treated with IgG-control or anti-PD1. IgG n=8 mice. Anti-PD1 n=10 mice. Mann-Whitney *U*-test. i) Quantification of tumour burden (mm^3^) at day 284 for DEN mice fed the ALIOS diet treated with IgG-control or anti-PD1. IgG n=15 mice. Anti-PD1 n=12 mice. Mann-Whitney *U*-test. j) Representative images of livers at day 284 for DEN mice fed either a control diet or the ALIOS diet treated with IgG-control or anti-PD1. IgG n=15 mice; Anti-PD1 n=12 mice. k) Survival plot in DEN/ALIOS mice treated with IgG-control or anti-PD1 (censored at day 165 post-treatment). IgG n=7 mice. Anti-PD1 n=8 mice. Log-rank (Mantel-Cox) test. l) Representative images of H&E-stained non-tumour livers and PCNA-stained tumours from DEN/ALIOS mice treated with IgG-control or anti-PD1 at day 284. IgG n=15 mice; Anti-PD1 n=12 mice. Scale bar = 100 µm. Dots in Fig. 1b, c, h, i represent individual mice.

Although elevated numbers of circulating neutrophils are associated with reduced HCC survival^17^, by contrast an enrichment of tumour associated neutrophils (TANs) is reported to correlate with improved survival^18^. This latter observation indicates a potential for TANs to influence the progression of HCC and raises the question of whether immunotherapy is influencing TANs (and vice versa). Ly6G^+^ neutrophils were found to be present in both tumour and non-tumour tissue of orthotopic-HCC mice and were significantly elevated in both compartments in the presence of NASH and remained high with anti-PD1 therapy (**Fig 2a** and **Supplementary Data Fig. 2a**). Increased numbers of TANs were also a feature in the DEN/ALIOS model and the increase reached significance with anti-PD1 treatment (**Fig. 2b** and **Supplementary Data Fig. 2a**). In addition, TANs were elevated in choline deficient-high fat diet (CD-HFD) spontaneous NASH-HCC model, and were retained with anti-PD1 therapy which is reported to also fail in this model^9^ (**Supplementary Data Fig. 2b, c**). Thus, we consistently observe TANs to accumulate in NASH-HCC, independent of the model examined, and they are retained in the tumour with anti-PD1 therapy. TANs display functional heterogeneity including anti- or pro-tumour phenotypes that impact on tumour growth^19^. Using transcriptomic profiling of tumour-isolated Ly6G^+^ cells, we determined the phenotype of TANs from DEN/ALIOS tumours. To account for environmentally-induced differences in gene expression^20^, we compared TANs with peripheral blood and liver neutrophils. DEGs with increased expression were enriched for process networks associated with inflammatory (e.g. *Nfkb1/Rel*, *Mapk8/Jnk1*, *Mapk9/Jnk2*, *Icam1*) and calcium (e.g. *Itpr1, Plcb1, Plcg1*) signaling (**Fig. 2c** **and Supplementary Data Fig. 2d**). Genes associated with a pro-tumour neutrophil phenotype, including *Csf1, Ccl3, Vegfa* and *Ptgs2*^19–21^ were also significantly up-regulated in TANs (**Supplementary Data Fig. 2e**).

**Figure 2.**
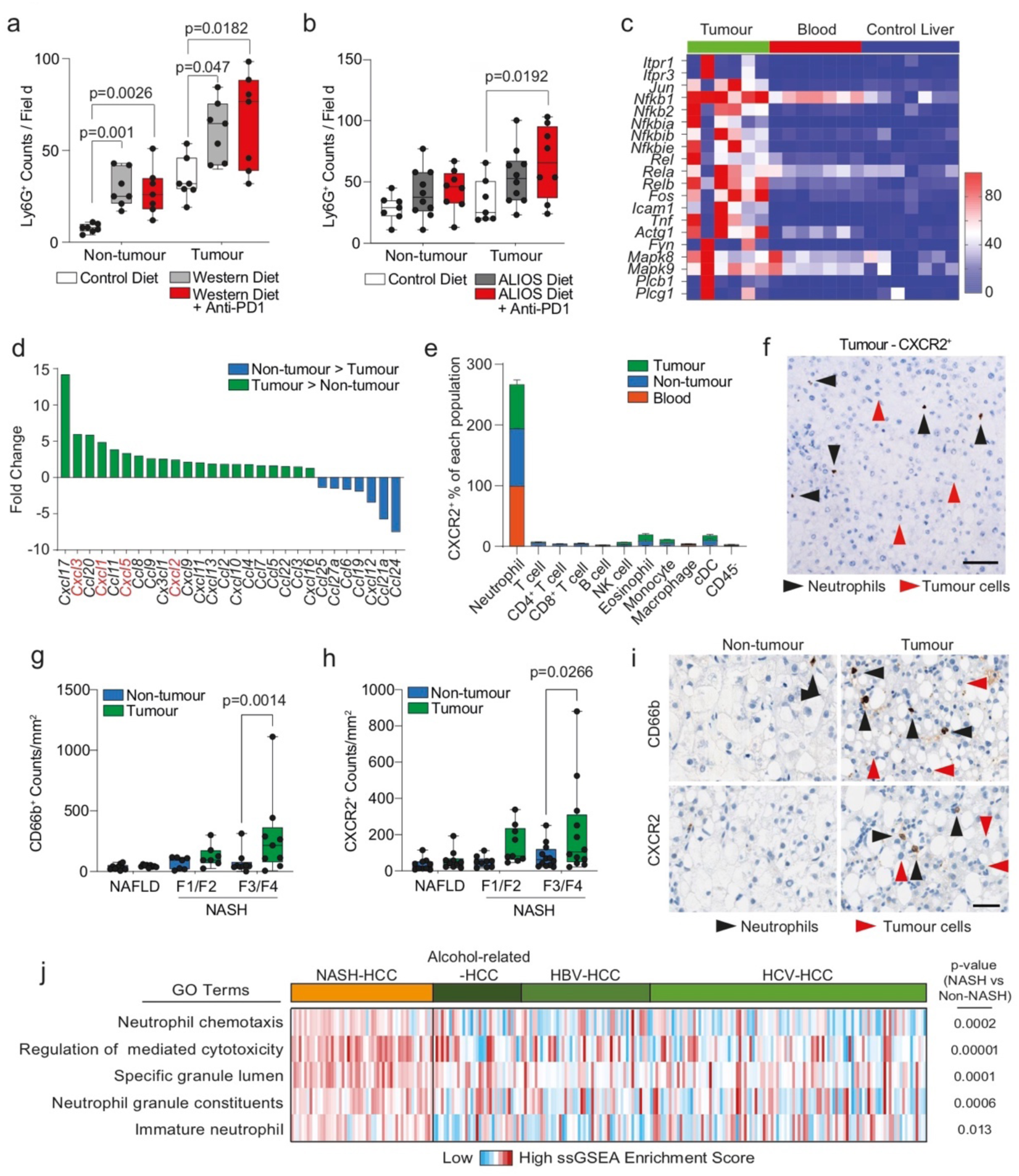
NASH-HCC and anti-PD1 resistance is associated with CXCR2^+^ neutrophils. a) Quantification of Ly6G^+^ counts/field in non-tumour liver and tumours from IgG-control or anti-PD1 treated orthotopic NASH-HCC mice. Control diet n=7, Western diet n=7, Western diet + Anti-PD1 n=7. Two-way ANOVA with Sidak’s multiple comparisons test performed. b) Quantification of Ly6G^+^ counts/field in non-tumour liver and tumours from IgG-control or anti-PD1 treated DEN/ALIOS mice. Control diet n=7, ALIOS diet n=10, ALIOS diet + Anti-PD1 n=8. Two-way ANOVA with Sidak’s multiple comparisons test performed. c) Heatmap showing row-scaled expression of DEGs associated with a pro-tumour neutrophil phenotype upregulated in DEN/ALIOS TANs compared with peripheral blood and control liver neutrophils. Data are from bulk Ly6G^+^ neutrophil RNA-Seq. d) Quantification of fold change for *Cxcl* and *Ccl* chemokine transcripts between DEN/ALIOS non-tumour liver and tumour. Data are from bulk tissue RNA-Seq. Non-tumour n=3 mice; tumour n=5 mice. e) Flow cytometric quantification of CXCR2^+^ as a percentage of cell populations in the peripheral blood, non-tumour liver and tumour in DEN/ALIOS mice. Peripheral blood n=5 mice; non-tumour liver n=5 mice; tumour n=3 mice. Error bars represent Mean ± SEM. f) Representative image of RNAscope *in situ* hydrisation staining of CXCR2 in DEN/ALIOS mouse tumours. Black arrowheads indicate positive infiltrating non-parenchymal cells and red arrows indicate negative tumour cells. g) Quantification of CD66b^+^ neutrophil counts/mm^2^ in non-tumour liver and tumour by IHC of non-alcoholic fatty liver disease (NAFLD)-HCC and NASH-HCC patient resected tissue. Patients were graded as NAFLD F0 (n=9), NASH F1/F2 (n=7), and NASH F3/F4 (n=9). Two-way ANOVA with Sidak’s multiple comparisons test. h) Quantification of CXCR2^+^ counts/mm^2^ in non-tumour liver and tumour by IHC from NAFLD-HCC and NASH-HCC patient resected tissue. Patients graded as NAFLD F0 n=10, NASH F1/F2 n=9 and NASH F3/F4 n=12. Two-Way ANOVA with Sidak’s multiple comparisons test. i) Representative IHC of CD66b^+^ and CXCR2^+^ cells (indicatd by arrowheads) in non-tumour and tumour regions of NASH-HCC patient-resected-tissue. Scale bar = 100 µm. n=12 patients. j) Heatmap showing row-scaled expression of neutrophil-associated process networks for human NASH-HCC compared with HBV, HCV and alcohol-related HCC (non-NASH-HCC). Data from bulk tumour microarray. Total n=237 patients. Dots in Fig. 2a, b, g, h represent individual mice.

Transcriptomic analysis of DEN/ALIOS tumours identified an up-regulation of myeloid associated cytokine and chemokine gene expression compared to normal liver (**Fig. 2d**). Notably, ligands (*Cxcl1, Cxcl2, Cxcl3, Cxcl5*) for the chemokine receptor CXCR2, the latter identified as being pre-dominantly expressed by Ly6G^+^ neutrophils, were all increased in tumour tissue (**Fig. 2d, e** and **Supplementary Data Fig. 2f**). *In situ* hybridation analysis of *Cxcr2* expression in DEN/ALIOS mouse tumours confirmed expression of *Cxcr2* to be specifically associated with morphologically identified infiltrating neutrophils and absent in parenchymal and tumour cells (**Fig. 2f**). This identifies CXCR2 as a neutrophil chemokine receptor that could be targeted to manipulate TANs in models of HCC-NASH^14^. In humans, the CXCR2 ligands *CXCL1* and *CXCL8* were significantly upregulated in NASH-HCC compared to NASH (**Supplementary Data Fig. 2g**). Neutrophil chemotaxis/migratory gene ontology (GO) terms were enriched in advanced human NASH (F4 fibrosis) and numbers of hepatic CD66b^+^ neutrophils increased with severity of NASH (**Supplementary Data Fig. 2h, i**). Moreover, in HCC patient tissue, CD66b^+^ neutrophils and CXCR2^+^ cells predominantly localised to NASH-HCC tumours with expression of the two markers correlating, and furthermore being demonstrated to be co-localised at the cellular level (**Fig. 2g, h** and **Supplementary Data Fig. 2j, k**). Similar to the mouse models, CXCR2 expression was limited to infiltrating immune cells and was absent on tumour epithelium within HCC in patients (**Fig. 2i**). We additionally noted that neutrophil expression signatures were enriched in human NASH-HCC compared to hepatitis B virus (HBV-HCC), hepatitis C virus (HCV-HCC) and alcohol-related-HCC^22^ (**Fig. 2j**). Thus, tumour infiltration of CXCR2-expressing neutrophils is characteristic of both murine models and human NASH-HCC and associates with resistance to anti-PD1 therapy in experimental models of NASH-HCC^9^.

### CXCR2 antagonism re-sensitises NASH-HCC to immunotherapy

We next determined the effects of a CXCR2 small molecule inhibitor (AZD5069^23^) in experimental NASH-HCC either administered alone or in combination with anti-PD1. We hypothesised that AZD5069 would suppress hepatic neutrophil recruitment. This was confirmed in the context of DEN-induced acute liver damage (**Supplementary Data Fig. 3a-c**). We also observed no change in F4/80^+^ macrophages and CD3^+^ T cells (**Supplementary Data Fig. 3d, e**). These data are consistent with previous studies showing that in acute inflammatory settings CXCR2 inhibition selectively reduces neutrophil recruitment^23^.

Treatments using either or both AZD5069/anti-PD1 were then investigated for their ability to suppress tumour growth in the DEN-ALIOS model (**Fig. 3a**). Tumour burden at day 284 was reduced for AZD5069 monotherapy and with combined AZD5069/anti-PD1 treatment compared to vehicle and anti-PD1 monotherapy, however, no change in tumour number was identified suggesting a suppression of cancer progression (**Fig. 3b** and **Supplementary Data Fig. 3f, g**). Examination of tumours revealed reduced numbers of epithelial mitotic bodies and a lower tumour-stage grading for the AZD5069/anti-PD1 group compared with other treatment arms including AZD5069 monotherapy without significantly altering the underlying NASH pathology (**Fig. 3c-f** and **Supplementary Data Fig. 3h**). This is clinically relevant as a high mitotic index in human HCC is a predictor of shorter disease-specific survival^24^. It was therefore noteworthy that the combination of AZD5069/anti-PD1 improved survival relative to monotherapies (**Fig. 3g**) Importantly, the benefits of AZD5069/anti-PD1 therapy were recapitulated in the orthotopic NASH-HCC model (**Fig. 3h-j**). In contrast to the DEN/ALIOS model, a lack of therapeutic effect was observed with either AZD5069 or anti-PD1 monotherapy (**Fig. 3i** and **Supplementary Data Fig. 3j**). However, AZD5069/anti-PD1 combination therapy reduced tumour burden at day 28 and extended survival relative to vehicle control and monotherapies (**Fig. 3i, j** and **Supplementary Data Fig. 3i**). Notably, the treatments had no influence on steatosis or body weight (**Supplementary Data Fig. 3j, k**). AZD5069/anti-PD1 treated mice reached clinical endpoint later, at which point tumour burden was similar between treatment arms, this being consistent with suppression of tumour growth (**Supplementary Data Fig. 3l**). Hence, although CXCR2 antagonism alone delivered modest model-dependant anti-tumour benefit, similar to observations made in models of non-hepatic cancers^25–30^, we show that CXCR2 inhibition sensitises to anti-PD1 immunotherapy in models of NASH-HCC that are otherwise resistant to anti-PD1 monotherapy.

**Figure 3.**
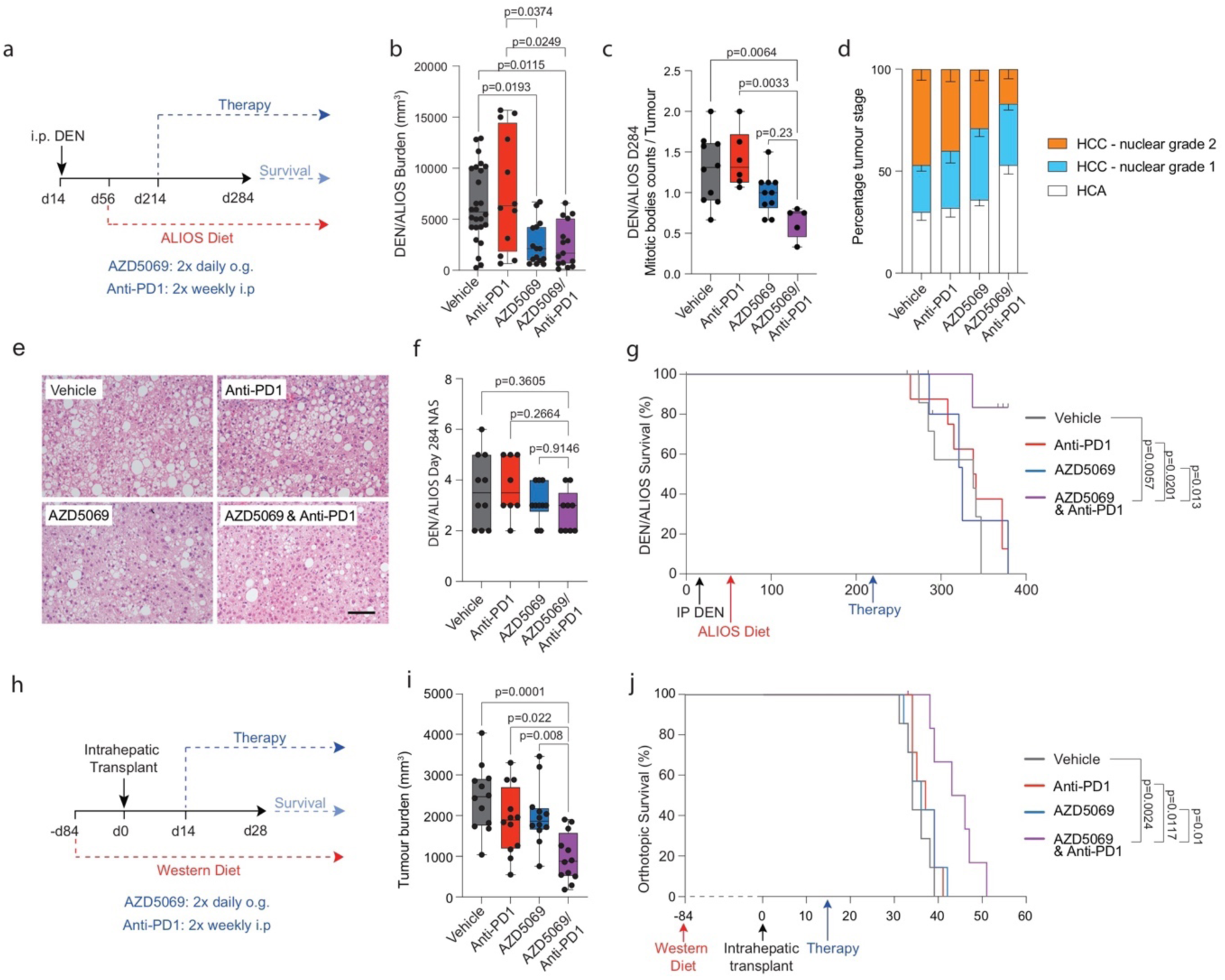
Inhibition of CXCR2^+^ pro-tumour neutrophils re-sensitizes NASH-HCC to anti-PD1 therapy. a) Schematic for DEN/ALIOS model treatment regime. b) Quantification of tumour burden (mm^3^) for DEN/ALIOS mice at day 284 for each treatment arm. Vehicle n=26 mice; anti-PD1 n=12 mice; AZD5069 n=15 mice; AZD5069/anti-PD1 n=15 mice. Kruskal-Wallis test with Dunn’s multiple comparisons test. c) Quantification of average mitotic body counts per tumour for DEN/ALIOS mice at day 284. All tumours analysed. Vehicle n=10 mice; anti-PD1 n=8 mice; AZD5069 n=10 mice; AZD5069/anti-PD1 n=9 mice. One-Way ANOVA with Tukey multiple comparisons test. d) Quantification of tumour stage based on nuclear grading for DEN/ALIOS mice at day 284 for each treatment arm. All tumours analysed. Vehicle n=10 mice; anti-PD1 n=8 mice; AZD5069 n=10 mice; AZD5069/anti-PD1 n=9 mice. Mean ± SEM. e) Representative images of non-tumour liver H&E for DEN/ALIOS mice. Vehicle n=7 mice; anti-PD1 n=8 mice; AZD5069 n=7 mice; AZD5069/anti-PD1 n=9 mice. Scale bar = 100 µm. f) Quantification of NAFLD activity score (NAS) in the livers for DEN/ALIOS mice at day 284. Vehicle n=10 mice; anti-PD1 n=8 mice; AZD5069 n=10 mice; AZD5069/anti-PD1 n=9 mice. g) Survival plot for DEN/ALIOS mice (censored at day 365). Vehicle n=7 mice; anti-PD1 n=8 mice; AZD5069 n=7 mice; AZD5069/anti-PD1 n=9 mice. Log-rank (Mantel-Cox) test. h) Schematic for orthotopic NASH-HCC model treatment regime. i) Quantification of tumour burden (mm^3^) for the orthotopic NASH-HCC mice at day 28. n=12 mice per condition. One-Way ANOVA with Tukey’s multiple comparisons test. j) Survival plot in orthotopic NASH-HCC mice. For each treatment group n=7 mice, 1 mouse censored due to non-liver related medical issue. Log-rank (Mantel-Cox) test. Dots in Fig. 3b, c, f, i represent individual mice.

### AZD5069/anti-PD1 therapy promotes an anti-tumour immune microenvironment

To further examine the concept that CXCR2 antagonism sensitises NASH-HCC to anti-PD1 therapy we asked if combination therapy activates classic T-cell mediated anti-tumour immunity. Characterisation of intratumoural T cells revealed intratumoural CD8^+^ T cells were significantly increased in both anti-PD1 and AZD5069/anti-PD1 therapy groups, with only anti-PD1 monotherapy significantly affecting CD4^+^ T cells (**Fig. 4a** and **Supplementary Data. Fig 4a)**. Combination therapy also enhanced intratumoural CD8^+^ T cell numbers in the orthotopic model (**Supplementary Data Fig. 4b**) Flow cytometric analysis revealed no gross phenotypic changes in early effector CD8^+^CD44^Hi^ T cells across treatment groups. However, anti-PD1 treatment alone significantly increased numbers of CD8^+^PD1^+^ T cells, this effect being recently reported by Pfister *et al*^9^ who suggested this T cell phenotype compromises the efficacy of anti-PD1 treatment in NASH-HCC (**Supplementary Data Fig. 4c**). The percentage of CD4^+^PD1^+^ T cells was also higher in the context of anti-PD1 monotherapy relative to other treatment groups (**Supplementary Data Fig. 4d**). Upregulation of Granzyme K (*Gzmk)*, and the T-box transcription factor Eomes combined with down-regulation of *Tbx21* is associated with an “exhausted-like” pro-inflammatory CD8^+^PD1^+^ T cell that has recently been reported to undergo clonal expansion during ageing^31^. Further investigation of T cell phenotype by RNAseq on isolated CD3^+^ cells revealed enhanced expression of *Gzmk* and *Eomes* following anti-PD1 therapy, both of which were suppressed when AZD5069 was combined with anti-PD1, whilst in contrast *Tbx21* expression was increased with combination therapy (**Fig. 4b**). Alongside these changes, AZD5069/anti-PD1 therapy enhanced the expression of Granzyme B (Gzmb), a cytotoxic serine protease expressed by neutrophils, NK cells and by recently activated CD8^+^ T cells and for which expression correlates with clinical outcome in PD1 immunotherapy^32–35^. Immunostaining of DEN/ALIOS tumours revealed that Gzmb was detected at low levels in vehicle and monotherapy groups yet in the context of AZD5069/anti-PD1 combination therapy was highly expressed and was localised within discrete immune cell clusters containing banded immature neutrophils and lymphocytes (**Fig. 4c** and **Supplementary Data Fig. 4e**). Enhanced Gzmb protein expression was also achieved with combination therapy in orthotopic tumours where we also noted that anti-PD1 monotherapy depressed expression of the protease relative to vehicle control (**Supplementary Data Fig. 4f**). These data led us to ask if depletion of CD8^+^ T cells would modulate the anti-tumour effects of AZD5069/anti-PD1 therapy. Depletion of CD8^+^ T cells was carried out by administration of anti-CD8*α* to mice bearing an orthotopic NASH-HCC tumour and alongside AZD5069/anti-PD1 treatment (**Fig. 4d**). Succesful depletion of CD8^+^ T cells was confirmed by an increase in the proportion of CD4^+^ cells relative to the total CD3^+^ population (**Supplementary Data Fig. 4g-i**) and resulted in a higher orthotopic tumour burden compared with IgG controls (**Fig. 4e**). The requirement for CD8^+^ T cells for the anti-tumour effect of combinaton therapy was additionally confirmed by performing anti-CD8*α*-mediated depletion in tumour-bearing DEN/ALIOS mice treated with combination AZD5069/anti-PD1 (**Supplementary Data Fig. 4j-m**).

**Figure 4.**
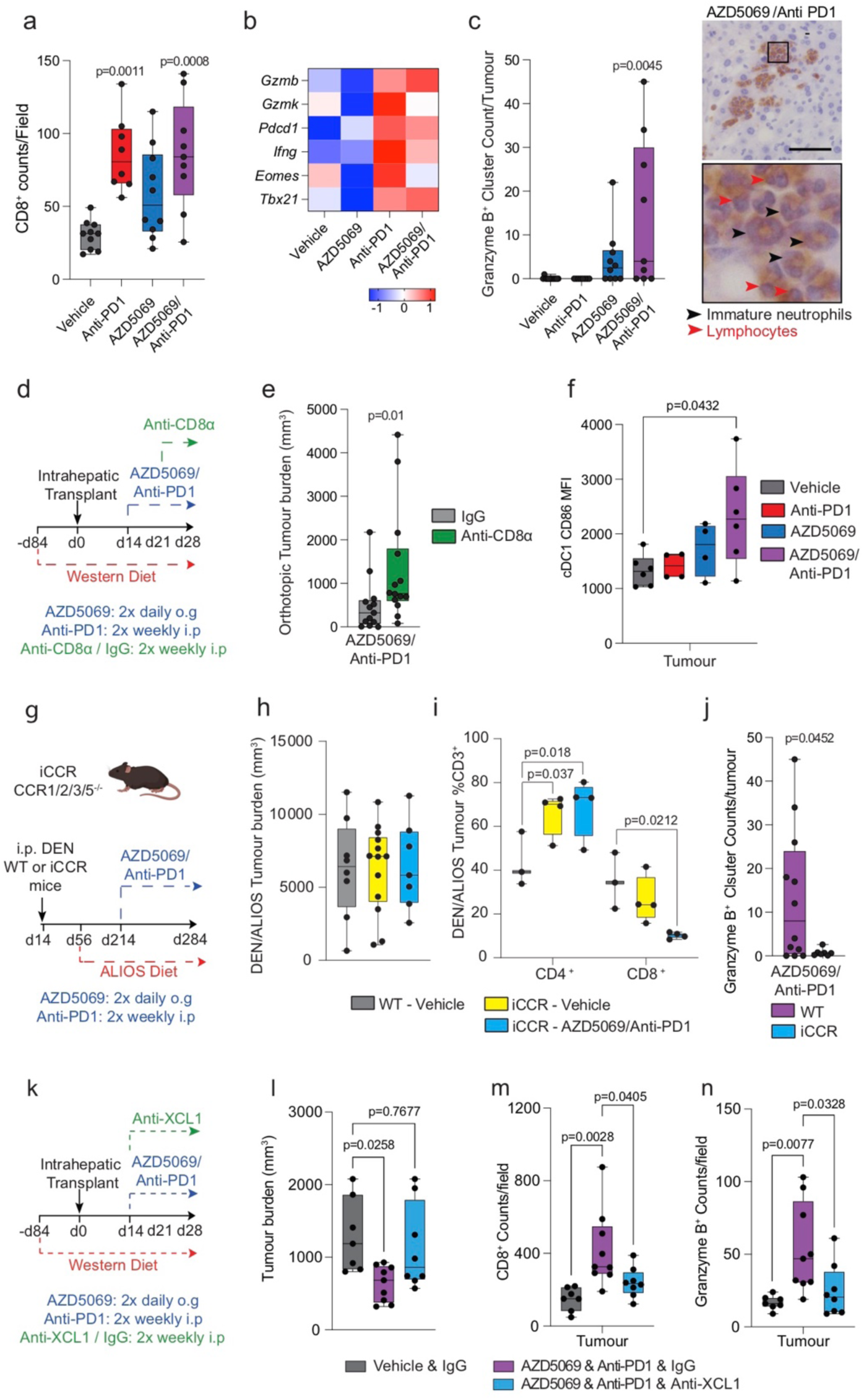
AZD5069/anti-PD1 therapy promotes an anti-tumour immune microenvironment. a) Quantification of CD8^+^ counts/field in tumours DEN/ALIOS model from each treatment arm. Vehicle n=10 mice; anti-PD1 n=8 mice; AZD5069 n=10 mice; AZD5069/anti-PD1 n=9 mice. One-Way ANOVA with Tukey multiple comparisons test. b) Heatmap showing row-scaled expression of genes associated with CD8^+^ T cell activation and exhaustion for DEN/ALIOS treatment groups. Data are from bulk CD3^+^ Tumour associated T cells analysed by RNA-Seq. c) Quantification of granzyme B^+^ clusters by IHC in the tumours for DEN/ALIOS mice from each treatment arm at day 284 and representative images of granzyme B^+^ clusters in AZD5069/anti-PD1 treated mice (black arrow heads = banded neutrophils; blue arrow heads = lymphocytes). Vehicle n=12; anti-PD1 n=8; AZD5069 n=10; AZD5069/anti-PD1 n=9. One-Way ANOVA with Tukey’s multiple comparisons test. Scale bar = 100 µm. d) Timeline schematic for the anti-CD8a depletion regime in the orthotopic NASH-HCC model. e) Quantification of tumour burden (mm^3^) in orthotopic NASH-HCC mice treated with AZD5069/anti-PD1 and IgG-control or anti-CD8α at day 28 post-intrahepatic injection. IgG n=13. Anti-CD8α n=14. Mann-Whitney *U*-test. f) Flow cytometric quantification of CD86 median fluorescence intensity (MFI) of intratumoural XCR1^+^ cDC1 cells from DEN/ALIOS mice treatment arms at day 284. Vehicle n=6 mice; anti-PD1 n=4 mice; AZD5069 n=4 mice; AZD5069/anti-PD1 n=6 mice. One-way ANOVA with Tukey multiple comparisons test. g) Timeline schematic for the DEN/ALIOS regimen and targeted therapies in mice with compound deletion of *Ccr1, 2, 3, 5* knockout mice, designated iCCR. h) Quantification of tumour burden for DEN/ALIOS mice Vehicle-treated WT and iCCR, and AZD5069/anti-PD1 treated iCCR mice at day 284. WT-Vehicle n=8 mice; iCCR-Vehicle n=13 mice; iCCR-AZD5069/Anti-PD1 n=7 mice. i) Flow cytometric quantification of CD4^+^ and CD8^+^ cells as a percentage of CD3^+^ T cells in tumours from WT-Vehicle, iCCR-Vehicle and iCCR-AZD5069/anti-PD1 treated DEN/ALIOS mice at day 284. WT-Vehicle n=3 mice; iCCR-Vehicle n=4 mice; iCCR-AZD5069/anti-PD1 n=4 mice. Two-way ANOVA with Tukey’s multiple comparisons test. j) Quantification of granzyme B^+^ clusters by IHC in WT and iCCR DEN/ALIOS mice treated with AZD5069/Anti-PD1 at day 284. WT n=12 mice. iCCR n=7 mice. Unpaired T-test. k) Timeline schematic for the anti-XCL1 neutralisation regime in the orthotopic NASH-HCC model. a) Quantification of tumour burden (mm^3^) in orthotopic NASH-HCC mice treated with vehicle control and IgG-control or AZD5069/anti-PD1 and either IgG-control or anti-XCL1 at day 28 post-intrahepatic injection. Vehicle and IgG n=7. AZD5069/anti-PD1 and IgG-control n=9. AZD5069/anti-PD1 and anti-XCL1 n=8. One-Way ANOVA with Tukey’s multiple comparisons test. l) Quantification of CD8^+^ counts/field in tumours of orthotopic NASH-HCC mice treated with vehicle control and IgG-control or AZD5069/anti-PD1 and either IgG-control or anti-XCL1 at day 28 post-intrahepatic injection. Vehicle and IgG n=7. AZD5069/anti-PD1 and IgG-control n=9. AZD5069/anti-PD1 and anti-XCL1 n=8. One-Way ANOVA with Tukey’s multiple comparisons test. m) Quantification of granzyme B^+^ counts/field in tumours of orthotopic NASH-HCC mice treated with vehicle control and IgG-control or AZD5069/anti-PD1 and either IgG-control or anti-XCL1 at day 28 post-intrahepatic injection. Vehicle and IgG n=7. AZD5069/anti-PD1 and IgG-control n=9. AZD5069/anti-PD1 and anti-XCL1 n=8. One-Way ANOVA with Tukey’s multiple comparisons test. Dots in Fig. 4a, c, e, f, h-j, l-n represent individual mice.

As recruitment and activation of XCR1^+^ conventional dendritic cells (cDC1) in tumours is considered critical for activation of cytotoxic CD8^+^ T cells and immunotherapy^36^ we next assessed CD86 surface expression as a marker of cDC1 activation in mice treated with AZD5069/anti-PD1 therapy. Anti-PD1 alone had no effect on activation of intratumour XCR1^+^ cDC1 cells compared to vehicle controls in the DEN/ALIOS model (**Fig. 4f**). AZD5069 alone also had no effect on activation of intratumour XCR1^+^ cDC1 cells, likely due to the limited expression of CXCR2 on cDCs (**Fig. 2e**). However, combined AZD5069/anti-PD1 therapy substantially increased the activation of intratumoural cDC1 cells (**Fig. 4f**). As several CC chemokines associated with DC recruitment were expressed in mouse NASH-HCC tumours responding to mono and dual therapies (**Supplementary Data Fig. 5a**) we next determine the effects of perturbing DC recruitment employing mice that are deficient for *Ccr1, Ccr2, Ccr3* and *Ccr5*, termed iCCR^37^. These mice were treated as per the DEN/ALIOS model, and AZD5069/anti-PD1 or control therapy administered (**Fig. 4g**). The number of cDC1 and cDC2 cells, and to a lesser extent F4/80^+^ macrophages but not neutrophils, were decreased in the tumours of iCCR mice (**Supplementary Data Fig. 5b-d**). Importantly, loss of myeloid recruitment alone in iCCR mice had no impact on tumour burden in the DEN/ALIOS model (**Fig. 4h**). However, unlike in wild-type (WT) mice, AZD5069/anti-PD1 therapy failed to reduce tumour burden in iCCR mice (**Fig. 4h**). This loss of effect was associated both with a reduction in tumour associated CD3^+^CD8^+^ T cells and loss of granzyme B^+^ immune clusters (**Fig. 4i, j**). To corroborate these data and to more specifically address the role of XCR1^+^ cDC1 cells we determined if AZD5069/anti-PD1 therapy of orthotopic NASH-HCC would be affected by anti-XCL1-mediated blockade of XCL1, a major chemokine involved in mediating cDC1 and CD8 T cell interactions (**Fig. 4k**)^38^. AZD5069/anti-PD1 therapy resulted in an increase in activated intratumoural XCR1^+^ cDC1 cells in line with observations in DEN/ALIOS mice, but with cDC1 activation being selectively suppressed upon treatment with anti-XCL1 (**Supplementary Data Fig. 5e, f**). This effect was associated with loss of the anti-tumoural action of AZD5069/anti-PD1 therapy; demonstrated by increased tumour burden in anti-XCL1 treated mice compared to IgG controls (**Fig. 4l**). Confirming an associated impact on cytotoxic T cells, AZD5069/anti-PD1-induced increases in tumoural cytotoxic CD8^+^ and GzmB^+^ cells which was suppressed when cDC1 activation was selectively blocked by anti-XCL1 (**Fig. 4m, n**). We conclude that combined suppression of CXCR2 and PD1 stimulates both tumoural recruitment and activation of cDC1 cells enabling T cell-mediated cytotoxicity.

### AZD5069/anti-PD1 therapy promotes tumour neutrophil accumulation and the formation of intratumoural immunological hubs

Given that CXCR2 is almost exclusively expressed on neutrophils (**Fig. 2e**) we were curious as to their role in AZD5069/anti-PD1 therapy and its associated tumour immune cell remodelling. Unexpectedly, we observed that combination therapy in both models of NASH-HCC was associated with a dramatic increase in TANs, whereas AZD5069 monotherapy brought about the anticipated reduction in TANs (**Fig. 5a, b** and **Supplemental Data Fig. 6a**). Real-time analysis of tumour neutrophil infiltration was not possible across the therapy time-course, so instead we examined circulating neutrophils sampled weekly from peripheral blood. Anti-PD1 alone had no demonstrable effects on circulating neutrophil numbers across the treatment period, whereas AZD5069 stimulated a transient increase in circulating Ly6G^+^ neutrophils peaking at 4 weeks after start of treatment (**Supplementary Data Fig. 6b**). A similar transient increase in circulating neutrophils was observed in AZD5069/anti-PD1 treated mice, however, this effect was delayed peaking at 6 weeks from start of treatment. These peripheral blood data indicated a change in neutrophil behaviour in response to dual long-term targeting of CXCR2 and PD1.

**Figure 5.**
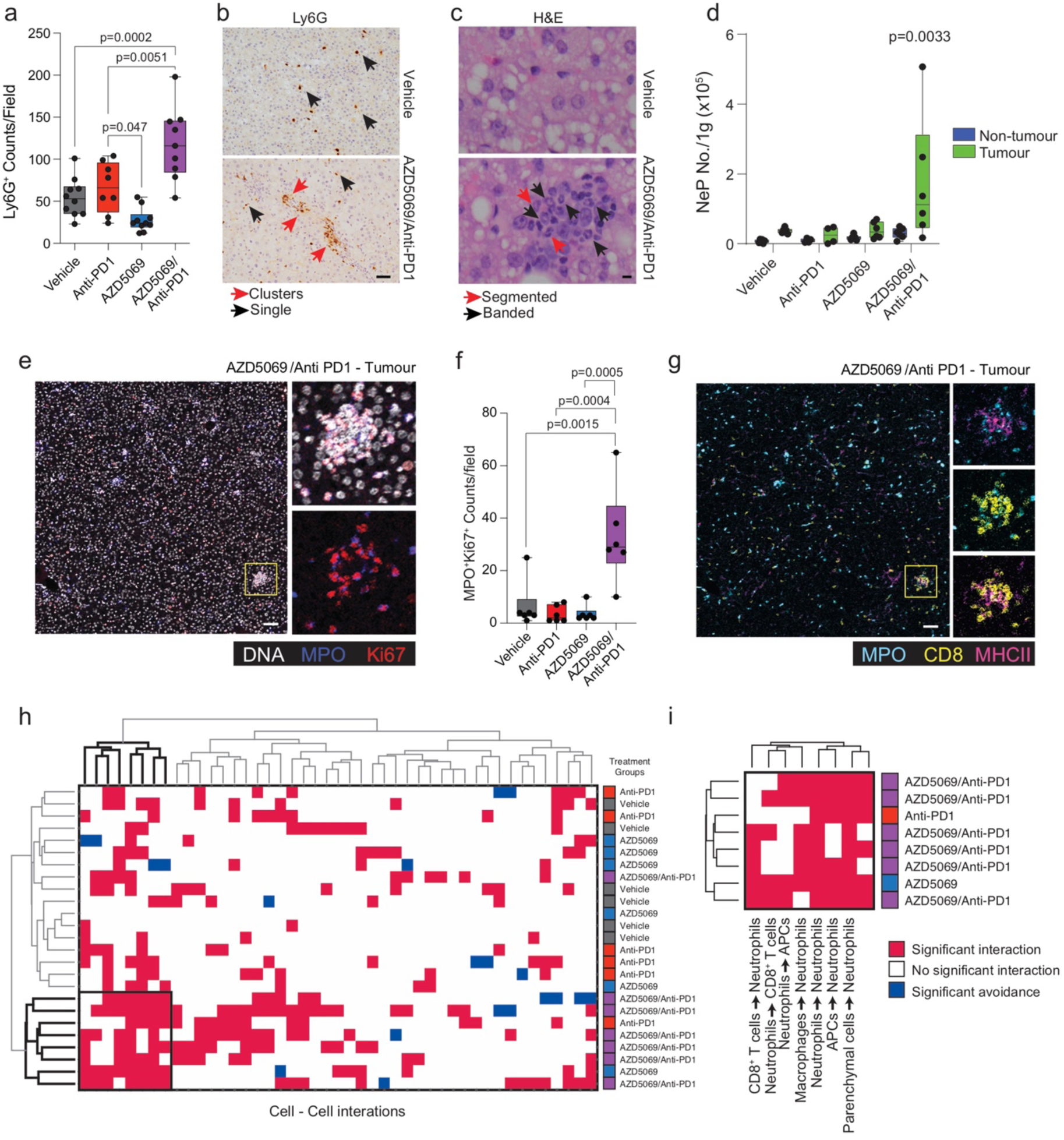
AZD5069/anti-PD1 therapy promotes tumour neutrophil accumulation and the formation of intratumoural immunological hubs. b) Quantification of Ly6G^+^ counts/field by IHC for DEN/ALIOS mice tumours at day 284. Vehicle n=10 mice; Anti-PD1 n=8 mice; AZD5069 n=10 mice; AZD5069/Anti-PD1 n=9 mice. One-Way ANOVA with Tukey’s multiple comparisons test. c) Representative images of Ly6G^+^ staining for DEN/ALIOS tumours. Vehicle n=10 mice; AZD5069/Anti-PD1 n=9 mice. Black arrows indicate single Ly6G^+^ neutrophils; red arrows indicate clusters of Ly6G^+^ neutrophils. Scale bar = 100 µm. d) Representative H&E from Vehicle-control and AZD5069/anti-PD1 treated mouse tumours identifying clusters of neutrophils with banded (blue arrows) and segmented (black arrows) nuclear morphology. Vehicle n=10 mice; AZD5069/Anti-PD1 n=9 mice. Scale bar = 10 µm. e) Flow cytometric quantification of NeP count/gram in non-tumour liver and tumour tissues for DEN/ALIOS mice for each treatment arm at day 284. Vehicle n=6 mice; anti-PD1 n=4 mice; AZD5069 n=6 mice; AZD5069/anti-PD1 n=6 mice. Two-way ANOVA with Sidak’s multiple comparisons test. f) Representative intra-tumour IMC image for DEN/ALIOS mice treated with AZD5069/anti-PD1. DNA = white; MPO = blue; Ki-67 = red. n=6 mice. Scale bar = 100 µm. n=6 tumours. g) Quantification of MPO^+^Ki-67^+^ counts/field for DEN/ALIOS tumours from IMC analysis. n=6 mice per condition. One-Way ANOVA with Tukey’s multiple comparisons test. h) Representative intra-tumour IMC image for DEN/ALIOS mice treated with AZD5069/anti-PD1. MPO = cyan; CD3 = yellow; MHCII = purple. n=6 mice. Scale bar = 100 µm. i) HistoCAT neighbourhood clustering analysis performed using phonograph clustered cell populations across all four treatment arms where red indicated a significant interaction, blue indicates a significant avoidance and white indicated no significant interaction. Each column represents the interaction of two cell types. Each row represents an individual mouse. Vehicle n=6 mice; anti-PD1 n=6 mice; AZD5069 n=6 mice; AZD5069/anti-PD1 n=7 mice. j) Magnified image of HistoCAT neighbourhood clustering analysis. Cluster showing specifically enriched cell-cell interactions. Cluster contains; anti-PD1 n=1 mouse; AZD5069 n=1 mouse; AZD5069/anti-PD1 n=6 mice. 7/8 cell-cell interactions characterised by antibodies used. Dots in Fig. 5a, d, f represent individual mice.

Immunohistochemical analysis of tumours identified clusters of TANs that were unique to AZD5069/anti-PD1 treatment and comprising a mixed population of banded and segmented neutrophil populations (**Fig. 5b, c** **and Supplementary Data Fig. 6c**). The presence of these clustered TANs in AZD5069/anti-PD1 treated HCCs was intriguing and suggestive of local proliferation. Zhu *et al*^39^ recently described early unipotent neutrophil progenitors (NeP) that produce neutrophils from adult BM. Conspicuously, NePs were significantly increased not only in the BM but also in tumours of AZD5069/anti-PD1 treated mice (**Fig. 5d** and **Supplementary Data Fig. 6d,e**). AZD5069/anti-PD1 treatment, therefore, alters granulopoiesis, while intratumour NePs may locally generate neutrophils, thus offering an explanation for the unexpectedly elevated numbers of TANs observed in mice receiving combination therapy.

To validate the presence of immature neutrophils in combined AZD5069/anti-PD1 treated tumours we utilised imaging mass cytometry (IMC) of tumour sections from DEN/ALIOS treatment arms (**Supplementary Data Fig. 6f-j)**. Neutrophils, both immature and mature, were identified as expressing the primary granule protein MPO. We confirmed intratumoural clusters of proliferating MPO^+^Ki67^+^ neutrophils to be significantly increased in AZD5069/anti-PD1 treated mice compared with monotherapies and vehicle controls (**Fig. 5e, f**). IMC neighbourhood analysis revealed intimate associations of MPO^+^Ki67^+^ neutrophils with CD8^+^ T cells and MHC Class II^+^ (MHCII^+^) antigen presenting cells (APCs) that were found in the regions of interest with six out of seven AZD5069/anti-PD1 treated tumours that were examined by IMC (**Fig. 5g-i**). In contrast, for anti-PD1 and AZD5069 monotherapies, IMC only detected these mixed immune cell hubs in one tumour for each type of treatment (**Fig. 5i**). Intravital microscopy confirmed the presence of stable tumour-associated Ly6G^+^ clusters, *in vivo*, in AZD5069/anti-PD1 treated mice (**Supplementary Data Fig. 6k, l)**. Directly interacting Ly6G^+^ TANs and CD3^+^CD8^+^ T cells that maintained physical contact over several minutes or more were also observed (**Supplementary Data Fig. 6m)**.

Longitudinal imaging of *ex vivo* precision cut liver slices (PCLS) was then performed to further interrogate Ly6G^+^ cell (neutrophil), CD8^+^ cell (T cell) and CD11c^+^ cell (DC and a subset of macrophages) dynamics within the tumours of DEN/ALIOS mice (**Supplementary Data Fig. 6n, o, Supplementary Movie)**. PCLS from AZD5069/anti-PD1 treated mice had the expected, elevated numbers of neutrophils, CD11c^+^ cells and CD8^+^ T cells (**Supplementary Data Fig. 6p-r)**. Although T cell speeds remained low in PCLS from all groups, neutrophil speed was increased in AZD5069/anti-PD1 treated tumours suggesting a more actively migrating phenotype for these neutrophils (**Supplementary Data Fig. 6s, t)**. Neutrophil-CD11c^+^ cell interactions were high in tumours irrespective of treatment, however, neutrophil-CD8^+^ T cell and CD11c^+^-CD8^+^ T cell interactions were elevated in AZD5069/anti-PD1 treated tumours compared to vehicle controls (**Supplementary Data Fig. 6u-y, Supplementary Movie).** These data provide evidence that combined therapeutic targeting of CXCR2^+^ neutrophils and the PD1-PDL1 immune checkpoint remodels the NASH-HCC tumour immune microenvironment, including the generation of locally proliferating immature neutrophil progenitors in close physical association with cytotoxic T cells.

### AZD5069/anti-PD1 combination therapy re-programmes the TAN phenotype

Given that our observations were consistent with intratumoural granulopoiesis in response to combination AZD5069/anti-PD1 therapy, we more closely characterised the TAN phenotype under these conditions. Grieshaber-Bouyer *et al*^40^, recently reported a chronologically ordered developmental path for neutrophils termed ‘neutrotime’. This extends from immature pre-neutrophils (early neutrotime) that are predominantly found in bone marrow (BM) to fully mature neutrophils (late neutrotime) mainly located in the circulation and spleen (**Supplemental Data Fig. 7a**). TAN transcriptome analysis revealed that AZD5069/anti-PD1 therapy induced neutrotime reprogramming along this neutrotime spectrum (**Fig. 6a** and **Supplementary Data Fig. 7b**). TANs in vehicle, anti-PD1 and AZD5069 treated tumours phenotypically resembled mature neutrophils, expressing genes characteristic of the late neutrotime (e.g. *Jund, Csf3r*, *Rps27*) (**Fig. 6a** and **Supplementary Data Fig. 7b**). However, late neutrotime genes were comprehensively downregulated in TANs from AZD5069/anti-PD1 treated mice, with a corresponding upregulation of transcripts characteristic of the early neutrotime (e.g. *Mmp8, Retnlg, Ltf, Lcn2, Camp, Chil3, Tuba1b*, *Fcnb*). Lactoferrin (Ltf) was of particular interest amongst the early neutrotime genes as its protein has well documented anti-cancer activities; including the activation of DCs and macrophages and enhancing the cytotoxic properties of natural killer cells^41–43^. Staining for Lactoferrin in DEN/ALIOS tumours was elevated in AZD5069/anti-PD1 treated mice where the protein was localised to the neutrophil-rich immune clusters that included banded immature neutrophils (**Fig. 6b,c**). These observations indicate a potential mechanism by which reprogrammed TANs may network with other immune cells to enact anti-tumoural effects.

**Figure 6.**
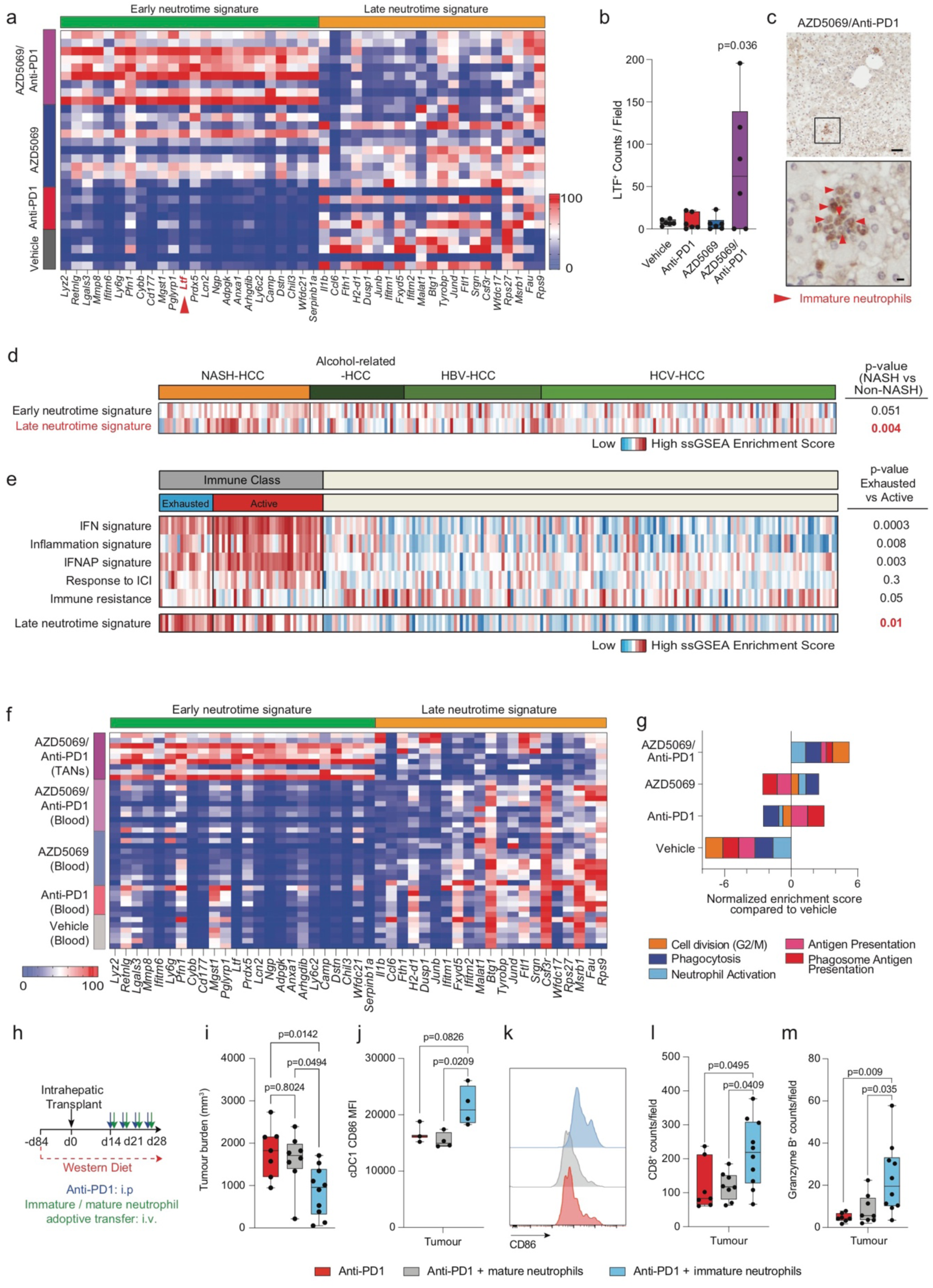
AZD5069/anti-PD1 combination therapy re-programmes the TAN phenotype. a) Heatmap showing row-scaled expression of genes associated with late and early neutrotime for DEN/ALIOS mice TANs. Data are from bulk Ly6G^+^ TANs analysed by RNA-Seq. b) Quantification of LTF^+^ counts/field by IHC for DEN/ALIOS mice tumours at day 284. Vehicle n=6 mice; Anti-PD1 n=6 mice; AZD5069 n=6 mice; AZD5069/Anti-PD1 n=6 mice. One-Way ANOVA with Tukey’s multiple comparisons test. c) Represenative image of LTF positive neutrophils (red arrow) in the tumour of AZD5069/anti-PD1 treated DEN/ALIOS mice at day 284. Scale bar top = 100 µm, bottom = 10 µm. d) Heatmap showing row-scaled expression of early and late neutrotime signatures for human NASH-HCC compared with HBV, HCV and alcohol-related HCC (non-NASH-HCC). Data are from bulk tumour microarray. In total n=237 patients analysed. e) Heatmap showing row-scaled expression of published HCC immune class signatures; IFN, inflammation, IFNAP, Response to ICI and immune resistance as well as the late neutrotime signature for human HCC active and exhausted immune subsets. Data are from bulk tumour microarray. In total n=228 patients analysed. f) Heatmap showing row-scaled expression of genes associated with late and early neutrotime signatures (Grieshaber-Bouyer *et al.,* 2021) for DEN/ALIOS peripheral blood neutrophils and AZD5069/Anti-PD1 treated TANs. Data are from bulk Ly6G^+^ TANs analysed by RNA-Seq. g) Gene set enrichment analysis (GSEA) showing normalised enrichment scores for TAN process networks highly enriched in; Anti-PD1 vs Vehicle (Phagosome Antigen Presentation and Antigen Presentation), AZD5069 vs Vehicle (Neutrophil Activation and Phagocytosis), and AZD5069/Anti-PD1 vs Vehicle (G2-M). Data are from bulk Ly6G^+^ TANs analysed by RNA-Seq. h) Timeline schematic for neutrophil based therapy treatment regime in the orthotopic NASH-HCC model. i) Quantification of tumour burden (mm^3^) in orthotopic NASH-HCC mice treated with anti-PD1 and immature or mature neutrophils at day 28 post-intrahepatic injection. Anti-PD1 n=7. Anti-PD1/Mature neutrophils n=8, anti-PD1/Immature neutrophils n=10. One-way ANOVA with Tukey’s multiple comparisons test. j) Flow cytometric quantification of CD86 median fluorscent intensity (MFI) of intraturmoural XCR1^+^ cDC1 cells from orthotopic NASH-HCC neutrophil/anti-PD1 therapy mice at day 28. Anti-PD1 n=3. Anti-PD1/Mature neutrophils n=4, anti-PD1/Immature neutrophils n=4. One-way ANOVA with Tukey’s multiple comparisons test. k) Representative histogram plot of CD86 median fluorscence intensity (MFI) of intraturmoural XCR1^+^ cDC1 cells from orthotopic NASH-HCC neutrophil/anti-PD1 therapy mice at day 28. Anti-PD1 n=3. Anti-PD1/Mature neutrophils n=4, anti-PD1/Immature neutrophils n=4. l) Quantification of intratumoural CD8^+^ counts/field in the tumours of orthotopic NASH-HCC neutrophil/anti-PD1 therapy mice at day 28. Anti-PD1 n=7. Anti-PD1/Mature neutrophils n=8, anti-PD1/Immature neutrophils n=10. One-way ANOVA with Tukey’s multiple comparisons test. m) Quantification of Granzyme B^+^ counts/field in the tumours of orthotopic NASH-HCC neutrophil/anti-PD1 therapy mice at day 28. Anti-PD1 n=7. Anti-PD1/Mature neutrophils n=8, anti-PD1/Immature neutrophils n=10. One-way ANOVA with Tukey’s multiple comparisons test. Bulk DEN/ALIOS Ly6G^+^ TAN RNA-Seq data in Fig. 6b, e, f: Vehicle n=6 mice; anti-PD1 n=5 mice, AZD5069 n=10 mice; AZD5069 n=8 mice. Dots in Fig. 6b, i, j, l, m represent individual mice.

Interrogation of transcriptome data from NASH- and non-NASH-related HCC patients identified the late neutrotime signature to be significantly enriched in NASH-HCC when compared to HCCs of other aetiologies^44^ (**Fig. 6d**). Moreover, the late neutrotime profile was specifically associated with human HCCs that are stratified by gene expression to the immune class and specifically within this group to the exhausted immune class which are typically resistant to immunotherapy (**Fig. 6e**)^44–47^. This suggests that TANs in human NASH-HCC resemble the mature phenotype of TANs in mouse NASH-HCC and may play a role in preventing ICI responses in patients, and as such we speculate they may be susceptible to similar therapeutic neutrotime reprogramming with AZD5069/anti-PD1 treatment. To examine the stage at which neutrophils are reprogrammed we compared tumoural to circulating neutrophil profiles in the treatment groups in the DEN/ALIOS model. Neutrotime reprogramming was specific to the intratumoural population of AZD5069/anti-PD1 treated mice and noteably without an associated neutrotime change in circulating neutrophils of this treatment group (**Fig. 6f**), this observation being consistent with tumour-selective neutrophil reprogramming. Hence, while the combination therapy brings about reprogramming of TAN maturity, it leaves intact the mature phenotype of circulating neutrophils required for their classic anti-microbial surveillance functions^48,49^.

RNA-Seq of purified Ly6G^+^ neutrophils revealed that TANs from AZD5069/anti-PD1 treatment mice were enriched for process networks associated with cell cycle, phagocytosis and antigen presentation when compared to vehicle controls (**Fig. 6g** and **Supplementary Data Fig. 7c**). AZD5069 monotherapy modestly enhanced the expression of signatures associated with cell division, phagocytosis, and degranulation while also eliciting a reduction in pro-tumour gene expression, with all of these effects being accentuated when AZD5069 was combined with anti-PD1 (**Fig. 6g** and **Supplementary Data Fig. 7c-e**). Anti-PD1 treatment promoted antigen presentation and processing signatures, which were also enriched in combined AZD5069/anti-PD1 treatment but not with AZD5069 monotherapy (**Fig. 6g**). These findings were again indicative of the combinatorial effects of AZD5069/anti-PD1 therapy on TAN phenotype. AZD5069 monotherapy (but not anti-PD1 monotherapy) suppressed the expression of key immune checkpoint molecules in TANs, including downregulation of *Cd80, Pvr, Sirpa, Pdl1* and *Pdl2*. This loss of immune checkpoint gene expression was maintained in the context of combination therapy and for some genes (e.g. *Pvr* and *Srpa*) we noted more pronounced suppressive effects when compared to the AZD5069 montherapy alone (**Supplementary Data Fig. 7f**). Hence, TAN-enriched immune hubs observed in AZD5069/anti-PD1 treated tumours are able to avoid immune checkpoint inhibition signals that might otherwise cause immune exhaustion. AZD5069/anti-PD1 TANs also displayed a strong correlation with transcriptional changes seen in neutrophils during an acute systemic inflammatory response^50^, including expression of genes involved in exocytosis, myeloid cell activation and degranulation (**Supplementary Data Fig. 7g, h**). Finally, these AZD5069/anti-PD1 TANs closely resembled a recently objectively characterised acute-inflammatory immature-Ly6G^Int^ neutrophil population isolated from lipopolysaccharide-(LPS)-treated mice^50^ (**Supplementary Data Fig. 7i**). In summary, AZD5069/anti-PD1 combination therapy brings about reprogramming of HCC-NASH TANs to exhibit immature, proliferative and inflammatory characteristics.

From these data we hypothesised that activated early neutrotime TANs have anti-tumoural properties. Due to their relatively low numbers and lack of specific surface markers it was not possible to isolate reprogrammed TANs from tumours in order to formally test this hypothesis. Instead, as proof-of-principe, we isolated inflammatory immature neutrophils enriched in the bone marrow of LPS-treated mice and a pool of mature bone marrow neutrophils isolated from control PBS treated mice. Adoptively transferring these cells to orthotopic NASH-HCC mice, we asked whether they would bring about an anti-tumoural effect in combination with anti-PD1 treatment (**Fig. 6h** and **Supplementary Data Fig. 7j, k**). Transfer of inflammatory immature neutrophils lead to a significant increase in circulating immature CXCR2^Lo^ neutrophils in the blood and resulted in a significant reduction in tumour burden (**Fig. 6i** and **Supplementary Data Fig. 7l**). In contrast transfer of mature neutrophils had no effect on tumour burden (**Fig. 6i**). To investigate underlying mechanism we examined intratumoural cDC1 and CD8^+^ T cells. Similar to treatment of mice with AZD5069/anti-PD1, we noted transfused immature neutrophils caused increased activation (CD86^+^) of intratumoural XCR1^+^ cDC1 cells and elevated CD8^+^ T cells in tumours, unlike mice transfused with equal numbers of mature neutrophils (**Fig. 6j-l**). Moreover, the adoptive transfer of neutrophils from LPS treated was associated with increased intratumoural Gzmb expression indicative of stimulation of cytotoxic activity within the tumour (**Fig. 6m**). Hence, we conclude that bone-marrow derived immature inflammatory neutrophils which have phenotypic similarities to AZD5069/anti-PD1 reprogrammed TANs are able to stimulate immune remodelling within HCC tumours and promote anti-tumoural effects.

## Discussion

Immune-based therapies hold considerable promise for the treatment of advanced HCC, however at present response rates are low and according to recent reports this is at least in-part determined by the immune cell composition of the tumour^45,51,52^. HCC on the background of NASH presents additional considerations because of the crosstalk occurring between inflammatory cells and various metabolic adaptions manifest in the disease such as insulin resistance, steatosis, oxidative stress and altered mitochondrial function^53^. Pfister and colleagues have reported that immunotherapy in NASH-HCC may be compromised due to high numbers of pro-tumour CD8^+^PD1^+^ T cells in the tumour microenvironment^9^. Here we show that selective targeting of neutrophils with a CXCR2 antagonist promotes the anti-tumour effects of anti-PD1 therapy in NASH-HCC, this effect being mechanistically associated with activation of classic CD8^+^ T cell and DC mediated anti-tumour immunity, but also with intratumoural reprogramming of TAN maturation and phenotype. Based on imaging mass cytometry we propose that the reprogrammed TANs, which are characterised by their proliferative and inflammatory characteristics, associate in tight clusters with CD8^+^ T cells and APCs to form anti-tumour Gzmb-secreting immune hubs within the NASH-HCC tumour microenvironment. Our work therefore emphasises the strong potential for targeted therapeutic manipulation of the innate immune system in cancer, but also uncovers a previously unrecognised crosstalk between the C-X-C chemokine/CXCR2 and PD1/PDL1 signaling systems that may be exploited to improve immunotherapy responses not only in NASH-HCC but also in other types of cancer that exhibit immunotherapy resistance^54,55^.

Neutrophil infiltration is a key pathological feature of human NASH that may result from upregulation of hepatic CXCL8 (IL-8) and CXCL1^56,57^, which we also report here to be enriched in human NASH-HCC. In addition, expression of CXCR2 on neutrophils in NASH is selectively enhanced through an auto-stimulation mechanism involving the upregulation of neutrophil-derived lipocalin 2^58^. Once present in the NASH and NASH-HCC microenvironments neutrophils are exposed to high levels of TGF-*β* which, as reported with other cancers^19–21^, can polarise TANs towards a so-called ‘N2’ tumour-promoting state^14^. It is also pertinent to address the relationship between TANs and myeloid-derived suppressor cells (MDSC), the latter being a heterogenous population comprising polymorphonuclear granulocytic Ly6G^+^Ly6C^Lo^ (PMN-MDSC) and monocytic Ly6G^-^Ly6C^Hi^ (M-MDSC) cells. Accumulating evidence suggests that PMN-MDSC are immunosuppressive neutrophils and may be functionally very similar to the TANs that have been termed ‘N2’, with shared pro-tumour properties^14,59^. In the mouse there are no markers to distinguish between PMN-MDSCs and neutrophils and as such we cannot rule out that TANs in mouse models of NASH-HCC include PMN-MDSCs which may also be susceptible to reprogramming in response to combined CXCR2 antagonism and anti-PD1 therapy. However, the typical inhibitory effects on DC and CD8^+^ T cell functions associated with the activities of PMN-MDSCs and immunosuppressive neutrophils were clearly overcome by combined AZD5069/anti-PD1 therapy.

A growing body of evidence suggests that CXCR2 inhibition may be therapeutically beneficial in many human cancers including; pancreatic, lung, ovarian, prostate, colon and now the liver^25–29,60^. Furthermore, in genetic murine models of lung cancer, inhibition of CXCR1 and 2 receptors in combination with anti-PD1 amplified anti-tumour responses^61,62^. The proposed mechanism of action, until now, however, was thought to rely on reprogramming of the tumour immune microenvironment, primarily as a result of impaired myeloid recruitment. The most remarkable immunobiological finding of our study was that, paradoxically, when combined with anti-PD1, CXCR2 inhibition leads to an increase in liver neutrophils and a selective reprogramming of the TAN neutrotime, with no evidence for a similar systemic effect on circulating neutrophils. The immature proliferative phenotype of the reprogrammed TANs evokes extramedullary granulopoiesis which can be seen in mice following antibody-mediated depletion of Ly6G^+^ cells and that is due to survival and expansion of residual tissue neutrophils driven by high systemic levels of granulocyte colony-stimulating factor^63^, indeed this rebound effect meant that we were unable to exploit this protocol to directly interrogate the function of reprogrammed TANs. However, as proof-of-principle we were able to establish that adoptive transfer of immature activated neutrophils isolated from bone marrow of LPS-treated mice has anti-tumour activity in NASH-HCC and this effect was accompanied by remodelling of tumour immunity including the activation of cDC1 cells, elevated CD8^+^ T cell counts and induction of anti-tumoural Gzmb; these being changes that were also noted with AZD506/anti-PD1 therapy. In future work it will be important to identify selective markers of the reprogrammed TANs that might be exploited for detailed functional characterisation, as well as for enabling their selective experimental manipulation which at present is not possible. Also, it will be important to determine precisely how and why combined CXCR2 antagonism and anti-PD1 treatment selectively induces proliferative immature neutrophils in the tumour. Clinically the ability to selectively reprogramme TANs while retaining mature anti-microbial neutrophils in the circulation may be very relevant in HCC since bacterial infections and septic shock are common clinical challenges in cirrhotic patients (in whom 90% of HCC develops)^64^.

In summary, we present a novel combination immunotherapy that enhances the efficacy of anti-PD1 in NASH-HCC. As the CXCR2 antagonist AZD5069 has been demonstrated to be safe for use in humans it is timely to determine if HCC patients would benefit from a similar therapy.

## Methods

### Mice, diets and treatments

All animal experiments using the orthotopic NASH-HCC model and DEN/ALIOS model were performed in accordance with a UK Home Office licence (PP8854860, PP390857 and PP0604995), adhered to ARRIVE guidelines (https://www.nc3rs.org.uk/arrive-guidelines), and in accordance with the UK Animal (Scientific Procedures) Act 1986, and were subject to review by the animal welfare and ethical review board of the University of Glasgow and Newcastle University. All mice were housed in specific pathogen free conditions with unrestricted access to food and water and maintained on a constant 12hr light-dark cycle under controlled climate (19-22 °C, 45-65% humidity). All animal experiments using the CD-HFD were performed in accordance with German law and the governmental bodies, and with approval from the Regierungspräsidium Karlsruhe (G11/16, G129/16, G7/17). Male mice were housed at the German Cancer Research Center (DKFZ) (constant temperature of 20-24 °C and 45-65% humidity with a 12hr light-dark cycle and were maintained under specific pathogen-free conditions.

For the orthotopic NASH-HCC model, intrahepatic injection of Hep-53.4 cells into the left lobe of C57BL/6 mice was performed under isoflurane general anaesthesia. Mice were fed *ad libitum* either a normal chow diet plus drinking water or a modified western diet (Envigo -TD.120528) plus sugar water (23.1 g/L fructose and 18.9 g/L glucose) for 3 months prior to implantation. For clinical relevance CXCR2smi and anti-PD1 therapeutic intervention was started at 14 days post implantation when small macroscopic tumours are present. Mice were then harvested at 28 days post implantation or left to reach an approved humane endpoint. To deplete CD8^+^ cells, mice received either anti-CD8a (Biolegend, 53-6.7) or IgG control (Biolegend, RTK2758; twice weekly i.p. 200μg) for 7 days after the initial 7 days of treatment with CXCR2smi and anti-PD1. Neutralization of XCL1 to deplete cDC1 cells; 50 μg of anti-XCL1 (R&D systems MAB486) or isotype-matched control antibodies (R&D systems MAB006) were injected i.p. twice a week for 2 weeks in combination with AZD5069/anti-PD1 therapy. Immature bone marrow neutrophil enrichment was performed with a single dose of lipopolysaccharide (LPS) from *E. coli*_0111:B4; (1 mg/kg, i.p.) or PBS control and femurs collected after 4 hours. Polymorphonuclear fraction of bone marrow from PBS and LPS treated mice isolated by percol density centrifugation. High density fraction, containing predominantly neutrophils was washed, resuspended, counted and 1×10^7^ cells i.v. injected twice weekly for 2 weeks into orthtopic NASH-HCC anti-PD1 treated mice. Tumour burden was calculated by measuring the size of the tumour in three perpendicular planes using digital callipers.

For the DEN/ALIOS model, WT C57BL/6 mice, bred in house were injected with a single dose of DEN at 80 mg/kg by i.p. injection at 14 days of age. Mice were placed on either the ALIOS diet consisting of an irradiated high trans-fat diet composed of 22% hydrogenated vegetable (Envigo, TD.110201) and sugar water or a control diet at 60 days of age. For clinical relevance, treatment with AZD5069 (AstraZeneca; 250 mg/mL in 0.5% Hydroxypropyl Methylcellulose (HPMC), 0.1% Tween 80; 100 mg/kg, o.g.) twice daily, or vehicle (0.5% HPMC, 0.1% Tween 80; o.g.) twice daily, anti-PD1 (Biolegend, RMP1-14; 200 µg, i.p.) bi-weekly or IgG (Biolegend, RTK2758; 200 µg, i.p.) bi-weekly was used to treat the mice. To deplete CD8^+^ cells in AZD5069/anti-PD1 DEN/ALIOS mice, mice received either anti-CD8a (Biolegend, 53-6.7) or IgG control (Biolegend, RTK2758; first dose i.p. 400μg and then twice weekly i.p. 200μg) from day 242 until harvest at day 284.

For the long-time CD-HFD feeding model, 5-week-old C57BL/6 mice (male) were fed choline-deficient high-fat diet (CD-HFD) (Research Diets; D05010402) for 13 months to induce NASH-HCC. For therapeutic intervention, anti-PD1 (BioXcell, RMP1-14; 200 µg, i.p.) bi-weekly, IgG (BioXcell, LTF-2; 200 µg, i.p.) bi-weekly, or vehicle (PBS; i.p.) bi-weekly, was used to treat the mice for 8 weeks.

For acute-DEN experiments, 12-15 week old WT C57B6/J mice bred in house or ordered in from Charles River were treated with a single dose of anti-PD1 or IgG control (dosing as above for DEN/ALIOS model) and/or AZD5069 or vehicle control (dosing as above for DEN/ALIOS model) twice daily. The following day mice received a single high-dose DEN injection (100 mg/kg, i.p.) and tissues were harvested the following morning.

For PBS-control and LPS-induced acute-inflammatory models, data is described in full in Mackey, *et al.,* 2021. Briefly, 8-10 week old WT C57BL/6 mice ordered in from Charles River were injected with a single dose of PBS (i.p) or LPS (*E. coli,* 0111:B4; 1 mg/kg, i.p.) and tissues were harvested after 24 hours.

### Human sample ethical approval

Collection and use of human tissue were ethically approved by the North East – Newcastle and North Tyneside 1 Research Ethics Committee. Human liver tissue from surgical resections was obtained under full ethical approval (H10/H0906/41) and through the CEPA biobank (17/NE/0070) and used subject to patients written consent. HCC tumour and non-tumour biopsy tissue was obtained under full ethical approval as approved by the Newcastle and North Tyneside Regional Ethics Committee, the Newcastle Academic Health Partners Bioresource (NAHPB) and the Newcastle upon Tyne NHS Foundation Trust Research and Development (R&D) department. (Reference numbers: 10/H0906/41; NAHPB Project 48; REC 12/NE/0395; R&D 6579; Human Tissue Act licence 12534).

### Scoring of tumour burden

For the DEN/ALIOS model, whole livers were weighed, then dissected into 3-4 sections and liver tumours were scored using digital callipers. For the orthotopic NASH-HCC model, whole livers were weighed and tumours were scored with digital callipers in three dimensions for calculating tumour volume (mm^3^).

### Sample processing and staining for flow cytometry and FACS

Tissues were collected in ice-cold PBS. Blood samples were collected into EDTA coated syringes and immediately treated with Erythrocyte Lysis Buffer containing NH_4_Cl, KHCO_3_ and Na_2_EDTA (Sigma-Aldrich) in dH_2_O, pH 7.2-7.4. Non-tumour liver and tumours were manually diced. Tumours were digested using a mouse tumour dissociation kit in GentleMACS C digestion tubes with a GentleMACS tissue dissociator (Miltenyi Biotec). Enzyme activity was neutralised by addition of cold RPMI/2% FBS and suspension was dispersed through a 70 µm cell strainer. Single cell suspensions were treated with RBC lysis buffer. Cells were blocked with CD16/32 (BioLegend) as required and stained with directly conjugated antibodies (listed below) for 25 minutes at 4 °C in the dark in PBS/1% BSA/0.05% NaN_3_. Zombie NIR (zNIR) fixable viability (1:1000; BioLegend) was added to exclude dead cells. For surface antigen staining only, cells were fixed in 2% paraformaldehyde. For intracellular staining, cells were fixed and permeabilized using the FOXP3/Transcription Factor staining buffer set (ThermoFisher), then staining of intracellular proteins. For cell counts, 10,000 AccuCount fluorescent particles (Spherotech) were added to each sample. All experiments were performed using a BD LSRFortessa flow cytometer using BD FACSDive^TM^ Diva software. Data were analysed using FlowJo software version 10.7.1.

All antibodies were purchased from BioLegend, except for CD101-PE, CD101-PE/Cy7, CD45-SB600 and IFNγ-PE/Cy7, which were obtained from eBioscience and Siglec-F-APC/Cy7, Siglec-F-Bv605, Ly6G-Buv395, CD11b-FITC, and CD162-Bv510, which were obtained from BD Biosciences.

General immune panel I: CXCR2-FITC (1:100, SA04E51), CD45-Bv711 (1:200, 30-F11), Siglec-F-Bv605 (1:200, E50-2440), CD31-PerCPCy5.5 (1:100, 390), CD3ε-Bv421 (1:100, 145-2C11), Ly6G-Buv395 (1:100, 1A8), CD8α-AF700 (1:100, 53-6.7), CD19-AF647 (1:100, 6D5), CD4-PE/Cy7 (1:200, RM4-5), Nkp46-PE (1:100, 29A1.4), fixable viability dye zNIR.

General immune panel II: CXCR2-FITC (1:100, SA04E51), XCR1-Bv785 (1:100, ZET), Ly6C-Bv711 (1:200, HK1.4), CD11b-Bv650 (1:400, M1/70), CD11c-Bv605 (1:200, N418), F4/80-Bv510 (1:200, BM8), CD115-Bv421 (1:200, AFS98), CD45-AF700 (1:200, 30-F11), CD172a-AF647 (1:100, P84), CD64-PE/Cy7 (1:200, X54-5/7.1), IA/IE-Bv421 (1:200, M5/114.15.2), CD26-PE (1:200, H194-112), fixable viability dye zNIR.

Tail bleed panel: Siglec-F-APC/Cy7 (1:200, E50-2440), CD45-Bv711 (1:200, 30-F11), CD62L-Buv395 (1:200, MEL-14), CD101-PE/Cy7 (1:200, Moushi101), Ly6G-AF647 (1:400, 1A8), CD11b-Bv650 (1:400, M1/70), CXCR2-FITC (1:100, SA04E51), CD3ε-Bv421 (1:100, 145-2C11), CD4-Bv605 (1:100, RM4-5), CD8α-AF700 (1:100, 53-6.7), PD-1-Bv785 (1:100, 29F.1A12), CD44-PerCP/Cy5.5 (1:100, IM7), PD-L1-PE (1:100, 10F.9G2), fixable viability dye zNIR.

Neutrophil panel: Siglec-F-APC/Cy7 (1:200, E50-2440), CD117-PE/Cy7 (1:100, 2B8), Ly6G-AF647 (1:400, 1A8), CD11b-Bv650 (1:400, M1/70), CXCR2-FITC (1:100, SA04E51), CD101 (1:200, Moushi101), CD45-Bv711 (1:200, 30-F11), PD-1-Bv785 (1:100, 29F.1A12), CD62L-Buv395 (1:200, MEL-14), fixable viability dye zNIR.

T cell panel: CD44-PerCP/Cy5.5 (1:100, IM7), CD4-FITC (1:100, RM4-5), PD-1-Bv785 (1:100, 29F.1A12), T-bet-Bv711 (1:200, 4B10), IL-17-Bv650 (1:400, TC11-18H10.1), CD45-SB600 (1:200, 30-F11), CD3ε-Bv421 (1:100, 145-2C11), CD19-Buv805 (1:400, 6D5), CD62L-Buv395 (1:200, MEL-14), CD8α-AF700 (1:100, 53-6.7), GranzymeB-AF647 (1:50, GB11), IFNγ-PE/Cy7 (1:200, XMG1.2), CD69-PE (1:100, H1.2F3) fixable viability dye zNIR.

DC and macrophage panel: CD45-AF700 (1:200, 30-F11), CD11c-Bv605 (1:200, N418), CD26-PE (1:200, H194-112), XCR1-Bv785 (1:100, ZET), CD103-PerCP/Cy5.5 (1:100, 2e7), IA/IE-Bv421 (1:200, M5/114.15.2), CD86-FITC (1:200, GL-1), CD11b-Bv650 (1:400, M1/70), CD172a-AF647 (1:100, P84), Ly6C-Bv711 (1:200, HK1.4), F4/80-Bv510 (1:200, BM8), CD64-PE/Cy7 (1:200, X54-5/7.1), fixable viability dye zNIR.

NeP panel: Siglec-F-Bv605 (1:200, E50-2440), FcεR1α-AF647 (1:400, MAR-1), CD16/32-PerCP/Cy5.5 (1:100, 93), Ly6B-FITC (1:400, 74), CD11a-PE (1:400, M17/4), Ly6G-PE/Cy7 (1:400, 1A8), CD162-Bv510 (1:400, 2PH1), CD115-Bv421 (1:200, AFS98), fixable viability dye zNIR.

Fluorescent-Activated Cell Sorting (FACS) panel: CD45-SB600 (1:200, 30-F11), CD48-PE/Cy7 (1:200, HM48-1), Ly6G-AF647 (1:400, 1A8), CD11b-FITC (1:400, M1/70), CD3ε-PE (1:100, 145-2C11), DAPI.

### Neutrophil and T cell RNA isolation and sequencing and analysis

Ly6G^+^ neutrophils and CD3^+^ T cells were FACS-sorted from the peripheral blood and tumours of DEN/ALIOS mice. Purity of isolated populations was analysed by flow cytometry at ≥ 97%. RNA was isolated using the Rneasy Micro Kit (Qiagen) according to the manufacturer’s protocol. RNA quality and quantity was checked on an Agilent Bioanalyser 2100 using RNA Pico 6000 chip. Libraries for cluster generation and DNA sequencing were prepared following the TaKaRa SMARTer Stranded Total RNA-Seq Kit-Pico Input Mammalian v2 protocol. Quality and quantity of the DNA libraries was assessed on an Agilent 2200 Tapestation (D1000 screentape) and Qubit (Thermo Fisher Scientific) respectively. The libraries were run on the Illumina Next Seq 500 using the High Output v2.5, 75 cycles kit (2 x 36cycles, paired-end reads). Illumina data were demultiplexed using bcl2fastq version 2.19.0, then adaptor sequences were removed using Cutadapt version 0.6.4 and quality checked using fastqc version 0.11.8. Next, paired end reads were aligned to the mouse genome version GRCm38.95 using HISAT2 version 2.1.0, and gene expression was determined using Htseq version 0.11.2. Differential expression analysis was performed using the R package DESeq2 version 1.22.2. Accurately identified genes were classed as those with ≥ 2 reads/million in ≥ half of the biological replicas in at least one experimental condition. DEGs were identified as those with a p-value ≤ 0.05 and fold change ≥ 1.5 between compared data sets. Gene ontology and pathway analysis was performed using MetaCore (Clarivate Analytics).

### Laser capture micro-dissection RNA-Sequencing

Formalin-fixed, paraffin-embedded (FFPE) 10μm liver sections on Zeiss membrane slides were dewaxed, rehydrated through graded alcohols, and then stained with haematoxylin and eosin (H&E). Tumour tissue was excised using Zeiss PALM microbeam laser capture microdissection microscope. RNA was then isolated using the High Pure FFPE RNA Micro Kit (Roche). RNA quality and quantity was checked using the DV200 metric (REF) on an Agilent Bioanalyser 2100 using RNA Pico 6000 chip. Total RNA sequencing libraries were prepared using the SMART-Seq Stranded kit [Takara Bio] following the manufacturer’s protocol. Libraries were quantified using and a Tapestation 4200 [Agilent] and Qubit 4 [Life Technologies] and equimolar pooled and sequenced at >30 million 100 bp single reads per sample on a NovaSeq 6000 using an 100 cycle SP flow cell [Illumina]. Data for individual samples was demultiplexed into separate FASTQ files using Illumina’s bcl2fastq software. Data were analysed as above (Neutrophil RNA isolation and sequencing and analysis).

### Whole tumour RNA-Sequencing and Analysis

Whole tumor and healthy tissue was snap frozen and stored at -80C. Tissue was homogenized using the Precellys Evolution homogenizer and bulk RNA was isolated using the Rneasy Kit (Qiagen) according to the manufacturer’s instructions, including the optional Dnase I step. RNA quality and quantity was analysed on a Nanodrop 2000 (Thermo Fisher Scientific) and an Agilent 2200 TapeStation (D1000 screentape). Only samples with a RIN value >7 were used for library preparation. Libraries were prepared using the TruSeq stranded mRNA Kit. Library quality and quantity were assessed using 2200 TapeStation (Agilent) and Qubit (ThermoFisher Scientific). The libraries were then run on an Illumina NextSeq 500 using the High Output 75 cycles kit (2×36cycle paired end reads). Data were analysed as above (Neutrophil RNA isolation and sequencing and analysis).

### RNA isolation, cDNA synthesis and qRT-PCR

Ly6G^+^ neutrophils sorted from orthtopic NASH-HCC tumours were snap frozen and stored at -80°c. RNA was isolated using the Rneasy Kit (Qiagen) according to the manufacturer’s instructions, including the optional Dnase I step. cDNA synthesis was performed using the iScript cDNA synthesis kit (Bio-Rad) according to the manufacturer’s instructions. Real time PCR was performed using SYBR Green jumpstart ready mix and the primers listed in (**Supplementary Table 1**).

### Publicly available gene expression dataset analysis

Publicly available RNA-seq datasets of differentially expressed genes from RNA-seq performed on biopsies from patients with NASH F0/F1 and NASH F4 were accessed. Gene ontology analysis was performed using genes significantly upregulated in patients with advanced disease. NASH-related HCCs, non-NASH-HCCs, as well as NASH liver human samples were previously described^22,44^. Transcriptomic data from human NASH liver (n=74) and NASH-HCC (n=53) samples were used to assess the expression of CXCR2 and key neutrophil-related cytokines in both tissues. The single sample Gene Set Enrichment Analysis (ssGSEA) module of GenePattern was used to determine the enrichment scores of immune-related gene signatures and signatures associated to response to immune checkpoint inhibitors^22,44^. Differentially expressed genes from RNA-seq performed on peripheral blood and liver Ly6G^+^ neutrophils, and peripheral blood Ly6G^Int^ and Ly6G^Hi^ neutrophils, from LPS-treated acute systemic inflammatory response and PBS-control treated mice were accessed. Gene ontology and pathway analysis was performed using MetaCore (Clarivate Analytics).

### Hep53.4 DNA isolation and analysis

DNA was isolated from Hep53.4 cells using the QIAamp DNA kit (Qiagen) as per manufacturers instruction and then sent for exome sequencing (Novogene). Raw read outputs were then passed through FastQC and FastQ screen for quality control. Raw reads were then aligned to the mm10 genome assembly using the Burrow-Wheeler aligner (BWA-MEM) software. Duplicate reads were identified using Picard tools and base recalibration was performed using BaseRecalibrator. Variants were called using Mutect2, before analysis using Ensembl Variant Effect Predictor to identify the likely effects of genomic variants. Data were visualised using R version 4.0.2 using the maftools, tidyverse packages and MutationalPatterns packages. We extracted known drivers of HCC in human disease from the Catalogue of Somatic Mutations in Cancer (COSMIC) database. Using mouse:human homology mapping through ENSEMBL cross species annotations, we then identified mutations present in known cancer driver genes. To account for gene length, the number of mutations is presented as number of mutations per million base pairs.

### Histology and immunohistochemistry

FFPE tissue sections were stained with H&E using established protocols. Immunohistochemistry (IHC) was performed on FFPE sections that were dewaxed and rehydrated through graded alcohols. Endogenous peroxidases were then blocked using hydrogen peroxide/methanol solution. Antigen retrieval was performed using Tris-EDTA pH9. Endogenous avidin and biotin were blocked using the Avidin/Biotin blocking kit (Vector Laboratories, SP-2001) and non-specific binding was blocked using 20% swine serum in PBS. Antibodies were incubated at room temperature for 1 hour. Sections were washed and incubated in the appropriate biotinylated secondary antibody. Sections were then washed and incubated in Vectastain Elite ABC-HRP reagent (Vector Laboratories, PK-6100). Staining was visualised using DAB substrate kite (Vector Laboratories, SK-4100), counter stained with mayer haematoxylin and then mounted. CD3 (AbCam Ab16669, pH6 1:50), CD4 (eBioscience 14-9766-82, ER2 Leica, 1:500), CD8 (eBioscience 14-0808-82, ER2 Leica, 1:500), Ly6G (clone 1A8, 2B Scientific BE0075-1, ER2 Leica, 1:60000), LTF (Thermofisher, PA5-95513, 1:200), granzyme B (Abcam, ab255598, Clone: EPR22645-206, 1:100), CD8 (Abcam, ab217344, Clone: EPR21769, 1:100). Image analysis was performed using a Nikon Eclipse Upright microscope and NIS-Elements BR analysis software. A minimum of ten consecutive non-overlapping fields were imaged where possible.

### Automated IHC and analysis of human tissue

Automated immunohistochemistry was performed using the Ventana Discovery XT platform. FFPE sections were dewaxed and rehydrated in EZ prep solution and then Tris-EDTA heat mediated antigen retrieval was performed. Endogenous peroxides and proteins were blocked using the Discovery Inhibitor CM. Sections were then incubated with primary antibodies (CD66b – Biolegend 305102, CXCR2 – Sigma HPA031999) followed by the appropriate secondaries. Staining was visualised by incubated slides in DAB followed by counterstaining with haematoxylin and then mounting. Sections were scanned using a Leica Aperio scanner and the analysis performed using Aperio ImageScope slide viewing software. Cell counts were performed on the whole tissue samples and normalised to the total area analysed (mm^2^).

### Imaging Mass Cytometry

The following antibodies were used for imaging mass cytometry: CK18 (Thermofisher, PA5-14263), F4/80 (Biorad, Cl:A3-1), CD3 (Biorad, CD3-12), Ki67 (Thermofisher, 14-5698-82), CD8 (Abcam, EPR21769), CD45R (Thermofisher, 14-0452-82), MHCII (Thermofisher, 14-5321-81), *α*SMA (ProteinTech, 14395-1-AP) and MPO (R&D, AF3667). Ly6G was not used to identify neutrophils as it was not compatible with IMC, instead MPO was used. Antibodies were first validated by immunohistochemistry and immunofluorescence staining of FFPE tissue samples using a single antigen retrieval method consisting of heat mediated antigen retrieval using the universal antigen retrieval solution (Abcam). Based on the relative immunofluorescence signal antibodies were ranked and then pair with the appropriate metal for conjugation to maximise signal to noise ratio. Antibodies were conjugated to metals using MaxPar antibody conjugation kits (Fluidigm) following manufacturer’s instructions. An antibody stabilisation solution was added before storage at 4°C (Candor Bioscience). The following metal-antibody conjugates were used: Nd145-CK18, Gd155-F4/80, Dy162-CD3, Dy163-Ki67. Dy164-CD8, Er166-CD45R, Tm169-MHCII, Er170-*α*SMA and Yb172-MPO. Conjugated antibodies were then validated signal by IMC in suspension mode (Helios) with antibody capture beads (AbC Total compensation beads, Thermo Fisher). To confirm conjugation did not alter antibody binding efficiency, antibodies were validated once more by immunofluorescence.

Tumour tissue microarrays were dewaxed and rehydrated through clearene and graded alcohols. Antigen retrieval was performed by immersing the slides in HIER Universal Antigen Retrieval reagent (Abcam, ab208572) at 95֯C for 20 min. Slides were then washed and blocked with 3% BSA in 1X PBS for 45 min. The cocktail of antibodies was prepared at the proper concentration of each one by diluting it in 0.5% BSA in 1X PBS and leaving it overnight at 4֯C in a humidified tray. The slides were then washed in 0.2% Triton X – 100 in PBS with gentle agitation, followed by rounds of 1X PBS. Nuclei were stained using DNA intercalator (Ir191/193) at 0.313μM in PBS for 30min. Finally, the slides were washed in ultra-pure water with gentle agitation and left air-drying at room temperature.

The Hyperion Tissue Imaging module was aligned and coupled to the Helios mass cytometry instrument and calibrated using the appropriate protocols (Fluidigm). Slide libraries were generated using low resolution images to aid identification of regions of interest (ROIs). ROIs (1mm^2^ in area) were ablated from each tumour sample on the TMAs. Imaging data files were exported in MCD viewer software (Fluidigm) as 16-bit single layer TIFFs. Single cell segmentation was performed using the Bodenmiller pipeline, combining open source software Ilastik for machine learning-based pixel classification and Cell Profiler for actual single cell segmentation. Cell phenotype and interaction analysis was performed in HistoCat.

### Liver intravital microscopy

Mice were anaesthetised and maintained using isoflurane in approximately 95% oxygen enriched air generated using a medical oxygen scavenger (VetTech). Mice were placed on a heat mat at 37 °C for the duration of the procedure. After loss of reflexes, the liver was exposed and a custom-built vacuum chamber fitted with a 13 mm glass cover slip placed onto the liver. Minimal suction (0.1-0.3 bar) was applied to stabilise the liver against the coverslip. Imaging was performed using an upright Zeiss LSM 880 Airyscan confocal microscope using a 20x/1 NA water immersion objective lens. Images were acquired in using a 32 channel Gallium arsenide phosphide spectral detector and signal was collected with a resolution of 8.9 nm over the visible spectrum. For visualisation of the vasculature and immune cell subtypes, fluorescently conjugated antibodies Ly6G (Biolegend, 5 µg, 1A8), CD45 (Biolegend, 5 µg, 30-F11), CD101 (eBioscience, 5 µg, Moushi101); CD3 (Biolegend, 5 µg, 145-2C11), CD8 (Biolegend, 5 µg, 53-6.7), CD31 (Biolegend, 10 µg, 390), were injected i.v. through the tail vein prior to anaesthesia. Livers were imaged for up to 60 minutes with a z-stack of 12 µm. At the end of the imaging session, mice were humanely killed by cervical dislocation under anaesthesia. Spectral images were unmixed with Zen software (Carl Zeiss) using references spectra acquired from unstained tissue (tissue autofluorescence) or slides labelled with individual fluorescently conjugated antibodies. Time-lapse images were visualized and analysed using IMARIS software (Bitplane, Oxford Instruments, Abingdon UK; v9). For interaction analysis, spots were initially automatically created for Ly6G^+^ cells and then corrected manually. Surfaces were initially automatically created for CD3^+^ cells and then corrected manually. Ly6G^+^ spots within 10μm, measured from the cell centre, of a CD3^+^ surface were considered interacting.

### Live precision cut tumour-containing liver slices (PCLS) imaging

Live PCLS procedure was adapted from McCowan et al for use with liver tissue^65^. Livers were sliced into 300μm thick sections on a vibratome and stained with Hoechst (1:2000) and directly conjugated fluorescent antibodies CD31 (Biolegend, 1:100, 390), CD8 (Biolegend, 1:100, 53-6.7), Ly6G (Novus bio, 1:100, 1A8), CD11c (Biolegend, 1:100, N418) and CD45 (Biolegend, 1:100, 30-F11) in complete medium (phenol-red free DMEM substituted with 1% FBS) for 20 minutes at 37°C. Slices were imaged on a Zeiss LSM880 confocal microscope in a full incubation chamber at 37°C with 5% CO2. Liver slices were imaged for 15-40min with z-stacks of 30µm. Acquisition was performed with a 32 channel Gallium arsenide phosphide(GaAsP) spectral detector using 20× objective. Samples were excited simultaneously with 405, 488, 561 and 633nm wavelength laser lines and signal was collected onto a linear array of the 32 GaAsp detectors in lambda mode with a resolution of 8.9 nm over the visible spectrum. Spectral images were then unmixed with Zen software (Carl Zeiss) using reference spectra acquired from unstained tissues (tissue autofluorescence) or beads labelled with single fluorophores.

### 4D images analysis

Timelapse images analysis and visualization was performed using Imaris (Bitplane). Neutrophils and T cells were detected and tracked using the ‘spot detection’ tool using either Ly6G^+^ fluorescence intensities (neutrophils) or CD8 (CD8^+^ T cells). CD11c^+^ areas were segmented using the ‘surface’ tool to measure total CD11c^+^ volume. All spots and surfaces were checked manually to avoid any false detections. Cell behavior was determined using the track speed (indicating cell mobility). Interacting cells were determined as cells located < 15µm to other spots or surfaces.

## Data Availability

All data will be deposited with accession codes, unique identifiers or web links for publicly available datasets provided before publication.

## Quantification and statistical analysis

Statistical analyses were performed using GraphPad Prism software (v9 GraphPad Software, La Jolla, CA, USA) and R (version 3.5.1) performing tests as indicated and were considered statistically significant. P-values are included in figures.

## Acknowledgements

**General:** We would like to that the core services and advanced technologies at the CRUK Beatson Institute with particular thanks to the Biological Services Unit, Molecular Services, Bioinformatics, Histology Department and the Beatson Advanced Imaging Resource. We would like to thank Danijela Heide, Jenny Hetzer for technical support for the CD-HFD NASH-HCC model. We would like to thank Catherine Winchester and Nathalie Sphyris for critical reading of the manuscript. We would like to thank the Newcastle University Comparative Biology Centre, Newcastle University Bioimaging Unit and the Newcastle University Flow cytometry core facility for technical assistance.

## Author Contributions

J.L., J.B.G.M., T.J. performed the majority of the laboratory-based work and analyses presented in the manuscript. E.R., T.M.D., F.F., W.C., K.G., A.H., C.N., S.L., M.L., R.P., P.K.H., R.E., C.E.W., M.R., C.F., D.F., S.R., N.W., M.M., A.C., D.G., A.F., D.M., A.F., X.C., N.M., C.A.F., X.L.R.I, A.J.M, M.M., R.R. performed a portion of the laboratory experiments and their related analyses. E.W.R., S.T.B., J.M.L., M.H., G.J.G. provided advice and/or contributed to the experimental design and writing. J.L., J.B.G.M., H.L.R., L.M.C., T.G.B., O.J.S., D.A.M. conceived the studies, designed the experiments and wrote the manuscript. All authors read and commented on the final manuscript. D.A.M., O.J.S., H.L.R., L.M.C., T.G.B. provided funding.

---

**Supplementary Data Figure 1.**
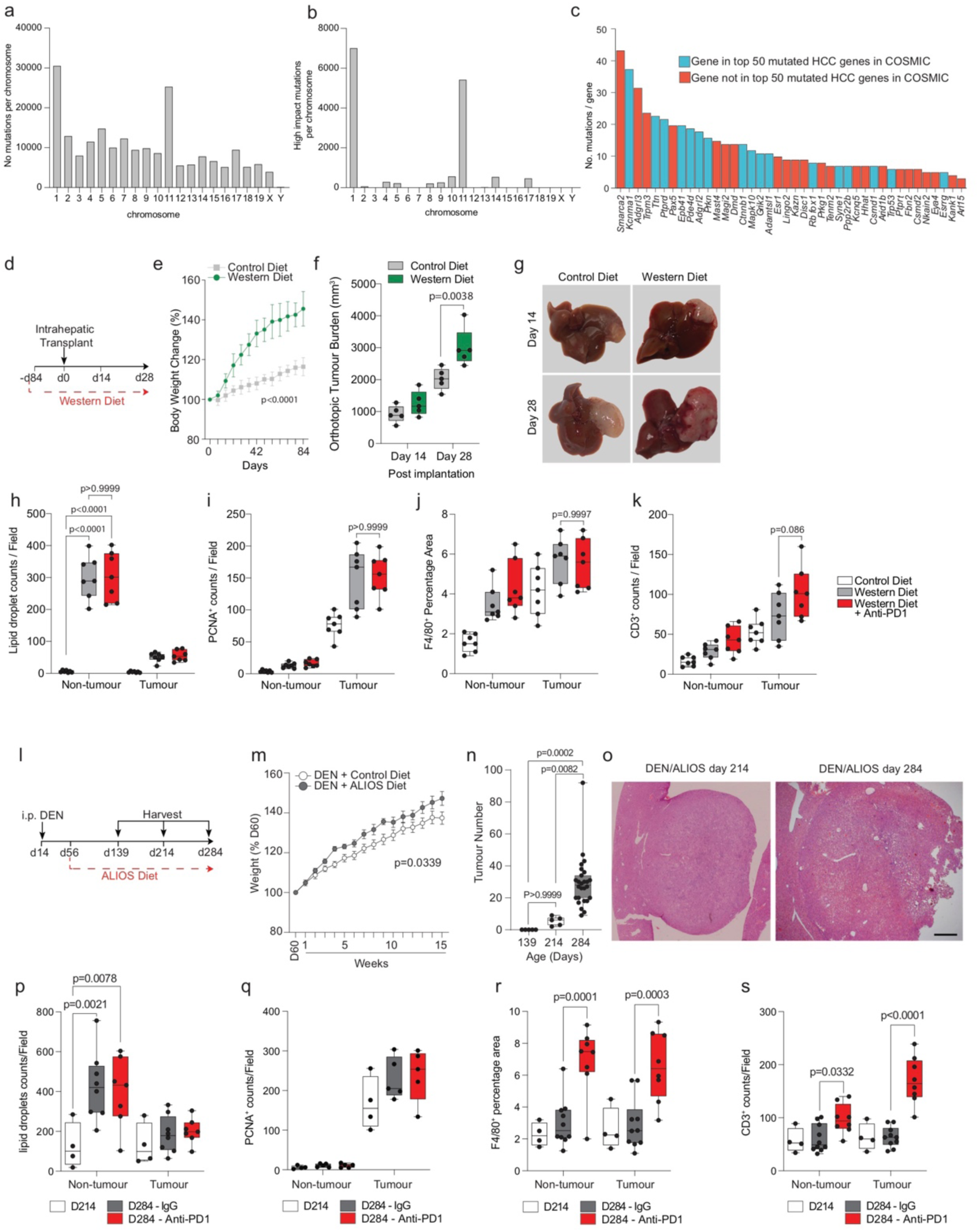
Steatosis promotes tumour development and anti-PD1 resistance. a) Number of mutations per chromosome in Hep53.4 HCC cells. b) Number of high impact mutations per chromosome in Hep53.4 cells. c) Top 40 mutated genes in Hep53.4 HCC cells, with high confidence human orthologues. Blue: gene in top 50 mutated HCC genes in the Catalogue of Somatic Mutations in Cancer (COSMIC) database. Red: gene not in top 50 mutated HCC genes in COSMIC. d) Timeline schematic of the orthotopic NASH-HCC model. e) Quantification of body weight change for mice fed a normal or Western diet, presented as percentage weight change compared with pre-diet weight. Control diet n=14 mice. Western diet n=14 mice. Two-way ANOVA. Error bars represent Mean ± SEM. f) Quantification of the tumour burden (mm^3^) for the orthotopic NASH-HCC mice fed a control or Western diet at day 14 and day 28 post intra-hepatic injection. n=5 mice per condition. Two-way ANOVA with Sidak’s multiple comparisons test. g) Representative images of livers at day 14 and day 28 post-intrahepatic injection for the orthotopic NASH-HCC mice fed a control or Western diet. n=5 mice per condition. h) Quantification of lipid droplet counts/field in non-tumour liver and tumour of the orthotopic NASH-HCC mice fed a control or Western diet and treated with IgG-control or anti-PD1, at day 28 post intrahepatic injection. n=7 mice per condition. Two-way ANOVA with Sidak’s multiple comparisons test. i) Quantification of PCNA counts/field in non-tumour liver and tumour for the orthotopic NASH-HCC mice fed a control or Western diet and treated with IgG-control or anti-PD1, at day 28 post intrahepatic injection. n=7 mice per condition. Two-way ANOVA with Sidak’s multiple comparisons test. j) Quantification of F4/80^+^ as a percentage of area in non-tumour liver and tumour for the orthotopic NASH-HCC mice fed a control or Western diet and treated with IgG-control or anti-PD1, at day 28 post intrahepatic injection. n=7 mice per condition. Two-way ANOVA with Sidak’s multiple comparisons test. k) Quantification of CD3^+^ counts/field in non-tumour liver and tumour for the orthotopic NASH-HCC mice fed a control or Western diet and treated with IgG-control or anti-PD1, at day 28 post intrahepatic injection. n=7 mice per condition. Two-way ANOVA with Sidak’s multiple comparisons test. l) Timeline schematic of the autochthonous DEN/ALIOS NASH-HCC model. m) Quantification of body weight change for mice injected with DEN at day 14 and fed a control or ALIOS diet from day 60, presented as a percentage of weight change compared with day 60. Control diet n=11 mice. ALIOS diet n=9 mice. Two-way ANOVA with Sidak’s multiple comparisons test. n) Quantification of tumour number for DEN/ALIOS mice at day 139, 214 and 284. Day 139 n=5 mice; day 214 n=5 mice; day 284 n=26 mice. Kruskal-Wallis test with Dunn’s multiple comparisons test. o) Representative image of H&E stained tumour nodules for DEN/ALIOS mice at day 214. n=5 mice. Scale bar = 1,000 µm. p) Quantification of lipid droplet count/field for DEN/ALIOS mice at day 214 and for DEN/ALIOS mice treated with IgG-control or anti-PD1 at day 284. Day 214 n=4 mice. Day 284 IgG n=8 mice. Day 284 anti-PD1 n=7 mice. Two-way ANOVA with Sidak’s multiple comparisons test. q) Quantification of PCNA^+^ as counts/field for DEN/ALIOS mice at day 214 and for DEN/ALIOS mice treated with IgG-control or anti-PD1 at day 284. Day 214 n=4 mice. IgG n=5 mice. Anti-PD1 n=5 mice. Two-way ANOVA with Sidak’s multiple comparisons test. r) Quantification of F4/80^+^ as percentage area for DEN/ALIOS mice at day 214 and for DEN/ALIOS mice treated with IgG-control or anti-PD1 at day 284. Day 214 n=4 mice. Day 284 IgG n=10 mice. Day 284 anti-PD1 n=8 mice. Two-way ANOVA with Sidak’s multiple comparisons test. s) Quantification of CD3^+^ count/field for DEN/ALIOS mice at day 214 and for DEN/ALIOS mice treated with IgG-control or anti-PD1 at day 284. Day 214 n=4 mice. Day 284 IgG n=10 mice. Day 284 anti-PD1 n=8 mice. Two-way ANOVA with Sidak’s multiple comparisons test. Dots in **Supplementary Data Fig. 1f, i-k, m, n, p-s** represent individual mice.

**Supplementary Data Figure 2.**
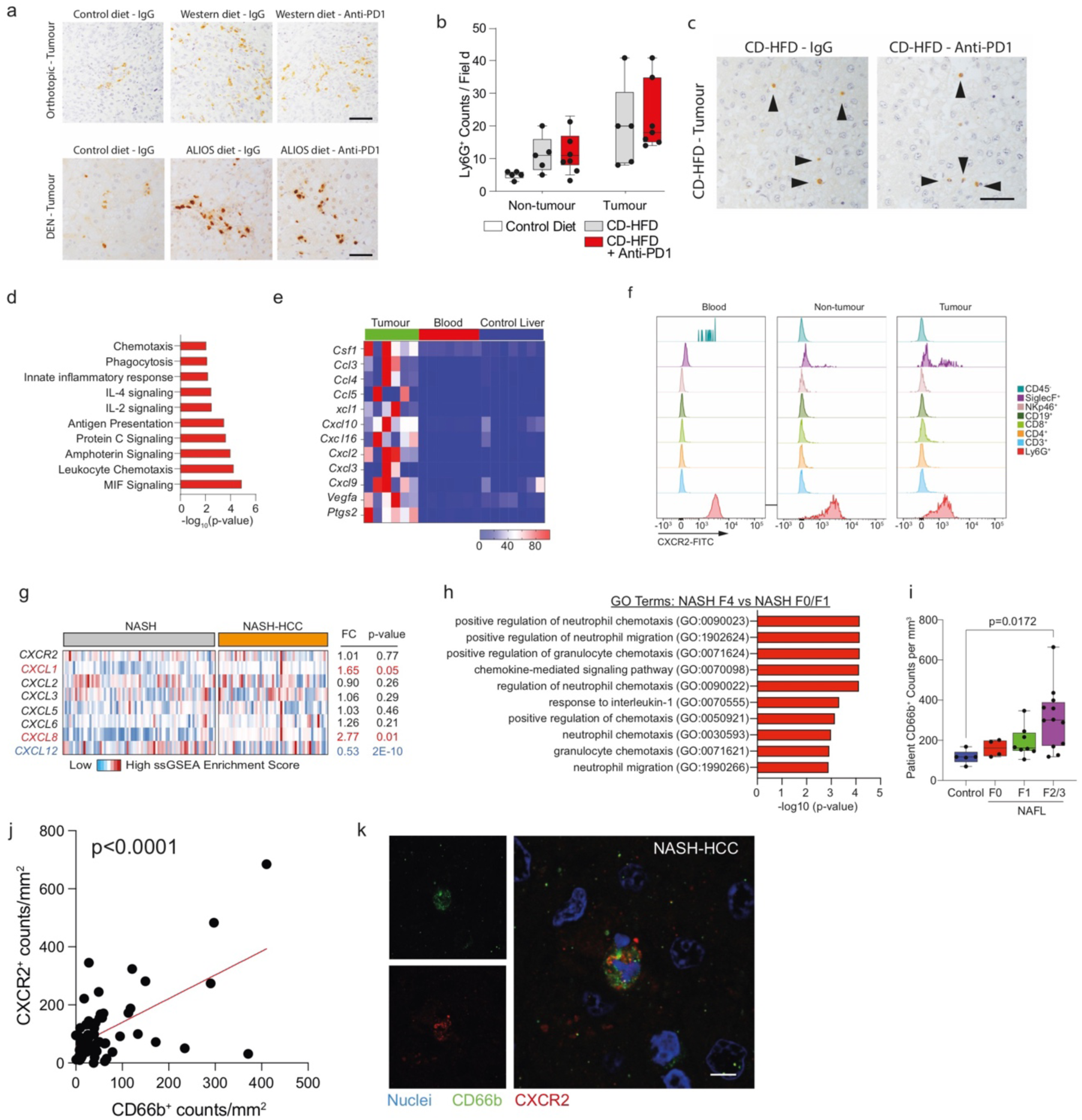
Neutrophils associate with NASH-HCC in mouse models and humans. a) Representative image of Ly6G^+^ neutrophil staining in tumours from the orthotopic NASH-HCC mice fed a control or western diet (top) and DEN mice fed a control or ALIOS diet (bottom). Orthotopic NASH-HCC: control diet + IgG-control n=7 mice; western diet + IgG-control n=7 mice; western diet + anti-PD1 n=7 mice. DEN/ALIOS: control diet + IgG-control n=7 mice; ALIOS diet + IgG-control n=10 mice; ALIOS diet + anti-PD1 n=8 mice. CD-HFD: IgG-control n=5 mice; CD-HFD + anti-PD1 n=7 mice. Scale bar = 100 µm. b) Quantification of Ly6G^+^ counts/field in non-tumour liver and tumours from IgG-control or anti-PD1 treated choline-deficient high fat diet (CD-HFD) mice. Non-tumour: control diet n=5 mice; CD-HFD n=5 mice; CD-HFD + anti-PD1 n=7 mice. Tumour: CD-HFD n=5, CD-HFD + Anti-PD1 n=7. One-Way ANOVA with Tukey multiple comparisons test. c) Representative images of Ly6G^+^ neutrophil staining in tumours from CD-HFD mice treated with IgG-control or anti-PD1. d) Top 10 process networks enriched in DEGs with increased expression in DEN/ALIOS TANs compared with peripheral blood and control liver Ly6G^+^ neutrophils. e) Heatmap showing row-scaled expression of pro-tumour associated neutrophil genes in DEN/ALIOS TANs compared with peripheral blood and control liver neutrophils. Data are from bulk Ly6G^+^ neutrophil RNA-Seq. f) Representative histograms for CXCR2 staining in immune cell populations isolated from the peripheral blood, non-tumour liver and tumours of DEN/ALIOS mice. n=3 mice. g) Heatmap showing row-scaled expression of *Cxcl* chemokine transcripts, related to neutrophil-function, for human NASH compared with NASH-HCC. Data are from bulk tumour RNA-Seq. NASH n=74 individuals; NASH-HCC n=53 individuals. h) GO Terms enriched in advanced (F4) compared with early stage (F0/F1) NASH patients. Data are from patient resected tissue RNA-Seq. Data accessed from Govaere *et al*. i) Quantification of CD66b^+^ neutrophil count by IHC from NAFLD patient resected tissue. Control n=5 individuals; F0 n=4 individuals; F1 n=8 individuals; F2/3 n=12 individuals. j) Correlation of CD66b and CXCR2 counts from tumour and non-tumour regions of HCC patients. k) Representative immunofluorescence of intra-tumour CXCR2^+^CD66b^+^ neutrophil from NASH-HCC patient resected tissue. Nuclei = blue; CD66b = green; CXCR2 = red. n=5 individuals. Scale bar = 10 µm. Dots/columns in **Supplementary Data Fig. 2e, g, i, j** represent individual mice/patients.

**Supplementary Data Figure 3.**
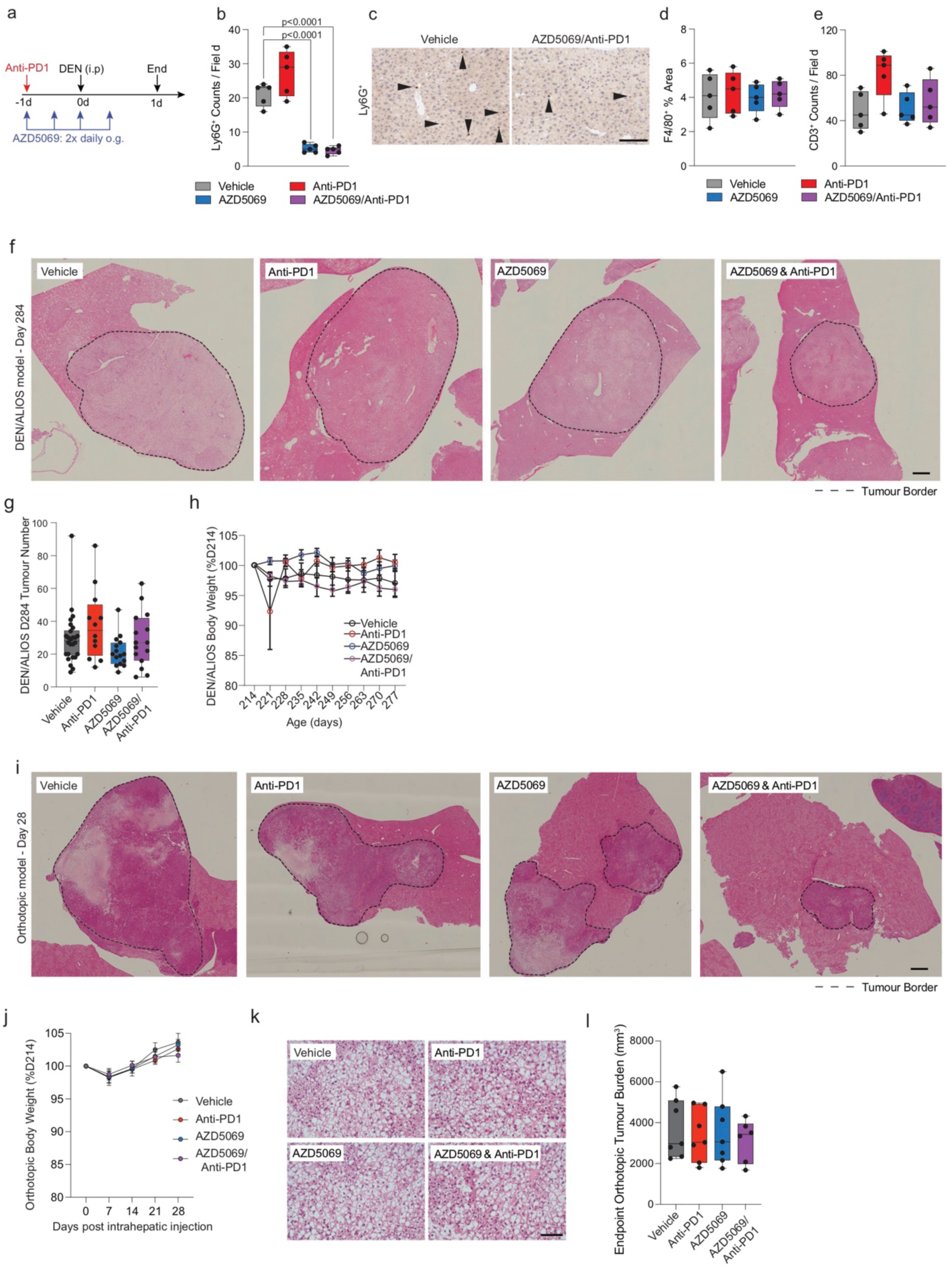
CXCR2 inhibition alters neutrophil regulation and sensitises to anti-PD1 therapy. a) Schematic for the acute-DEN model treatment regime. b) Quantification of Ly6G^+^ counts/field by IHC for the livers from acute-DEN mice treated with Vehicle-control, anti-PD1, AZD5069, or AZD5069/anti-PD1. n=5 mice per condition. One-Way ANOVA with Tukey’s multiple comparisons test. c) Representative Ly6G^+^ staining of liver sections from acute-DEN mice. n=5 mice per condition. Scale bar = 100 µm. d) Quantification of F4/80^+^ macrophages as a percentage area of the field by IHC for the livers from acute-DEN mice treated with Vehicle-control, anti-PD1, AZD5069, or AZD5069/anti-PD1. n=5 mice per condition. One-Way ANOVA with Tukey’s multiple comparisons test. e) Quantification of CD3^+^ counts/field by IHC for the livers from acute-DEN mice treated with Vehicle-control, anti-PD1, AZD5069, or AZD5069/anti-PD1. n=5 mice per condition. One-Way ANOVA with Tukey’s multiple comparisons test. f) Representative image of H&E stained liver sections for DEN/ALIOS mice at day 284. Vehicle n=15 mice; anti-PD1 n=12 mice; AZD5069 n=15 mice; AZD5069/anti-PD1 n=15 mice. Scale bar = 1,000 µm. g) Quantification of tumour number for DEN/ALIOS mice at day 284 for each treatment arm. Vehicle n=15 mice; anti-PD1 n=12 mice; AZD5069 n=15 mice; AZD5069/anti-PD1 n=15 mice. Kruskal-Wallis test with Dunn’s multiple comparisons test. h) Quantification of body weight change for DEN/ALIOS mice, presented as percentage of weight change compared with pre-treatment start (day 214). Vehicle n=16 mice; anti-PD1 n=14 mice; AZD5069 n=14 mice; AZD5069/anti-PD1 n=13 mice. Error bars represent Mean ± SEM. i) Representative image of H&E stained liver sections from the orthotopic NASH-HCC mice at day 28. n=12 mice per condition. Scale bar = 1,000 µm. j) Quantification of body weight change for the orthotopic NASH-HCC mice presented as a percentage of weight change compared with pre-intrahepatic injection. Error bars represent Mean ± SEM. n=12 mice per condition. k) Representative images of H&E stained livers from orthotopic NASH-HCC mice. Vehicle n=12, Anti-PD1 n=12, AZD5069 n=12 and AZD5069/Anti-PD1 n=12 mice. Scale bar = 100 µm. l) Quantification of tumour burden (mm^3^) at end-point for the orthotopic NASH-HCC mice. Vehicle n=7 mice; anti-PD1 n=7 mice; AZD5069 n=7 mice; AZD5069/anti-PD1 n=6 mice. Dots in **Supplementary Data Fig. 3b, c, d, g, I, m** represent individual mice.

**Supplementary Data Figure 4.**
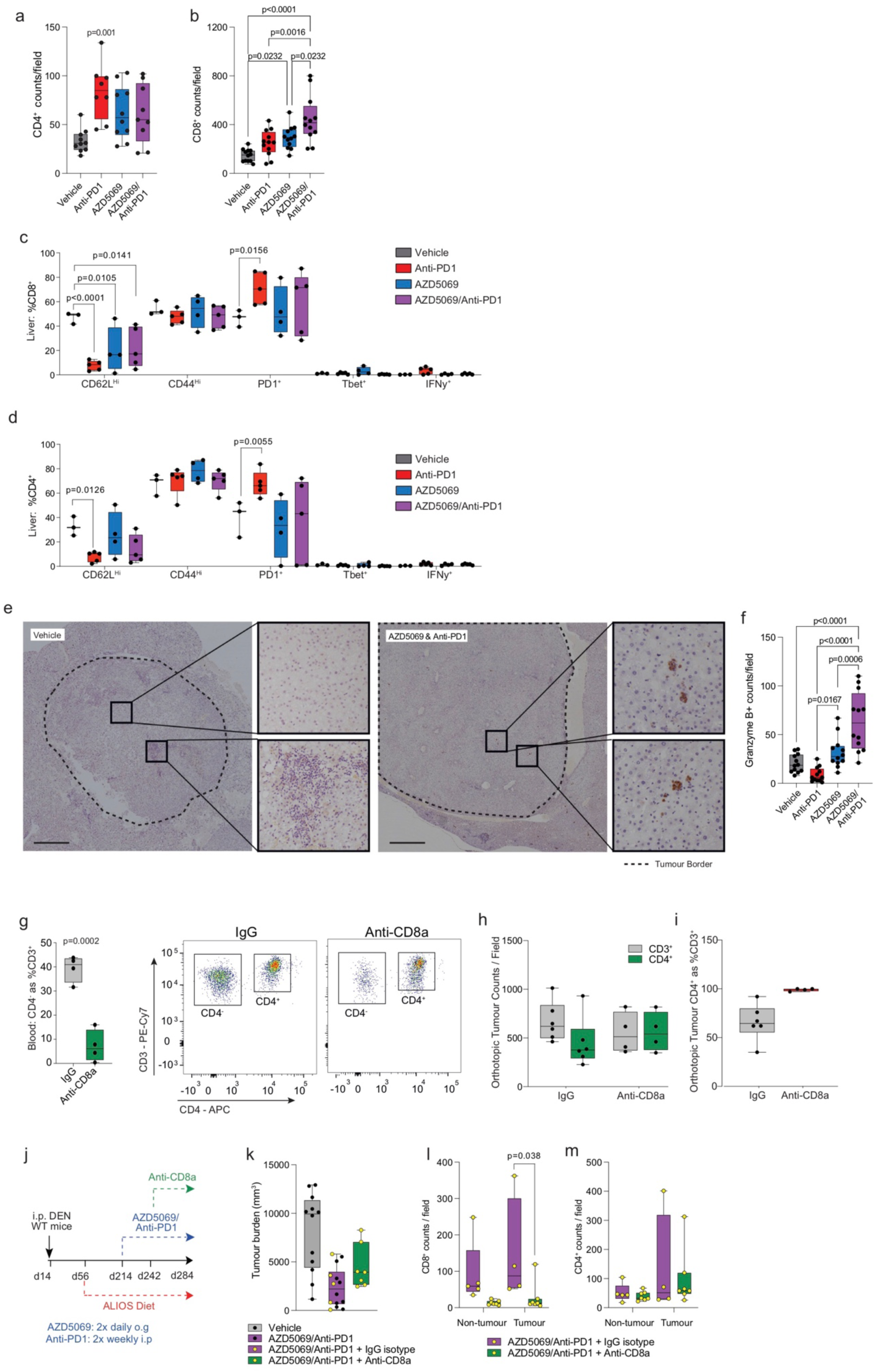
Cytotoxic CD8^+^ T cells contribute to combined AZD5069 & anti-PD1 anti-tumour phenotype. a) Quantification of CD8^+^ counts/field in tumours DEN/ALIOS model for each treatment arm. Vehicle n=10 mice; anti-PD1 n=8 mice; AZD5069 n=10 mice; AZD5069/anti-PD1 n=9 mice. One-Way ANOVA with Tukey multiple comparisons test. b) Quantification of CD8^+^ counts/field in livers for the orthotopic NASH-HCC mice for each treatment arm. Vehicle n=12 mice; anti-PD1 n=12 mice; AZD5069 n=12 mice; AZD5069/anti-PD1 n=12 mice. One-Way ANOVA with Tukey’s multiple comparisons test. c) Quantification of CD3^+^CD8^+^ cell surface phenotyping for DEN/ALIOS mice from each treatment arm at day 284. Vehicle n=3 mice; anti-PD1 n=5 mice; AZD5069 n=4 mice; AZD5069/anti-PD1 n=5 mice. Two-way ANOVA with Dunnett’s multiple comparisons test. d) Quantification of CD3^+^CD4^+^ cell surface phenotyping for DEN/ALIOS mice from each treatment arm at day 284. Vehicle n=3 mice; anti-PD1 n=5 mice; AZD5069 n=4 mice; AZD5069/anti-PD1 n=5 mice. Two-way ANOVA with Dunnett’s multiple comparisons test. e) Representative images of granzyme B^+^ clusters in Vehicle-control and AZD5069/anti-PD1 treated DEN/ALIOS livers. Scale bar = 1,000 µm. f) Quantification of Granzyme B^+^ counts/field in livers for the orthotopic NASH-HCC mice for each treatment arm. Vehicle n=12 mice; anti-PD1 n=12 mice; AZD5069 n=12 mice; AZD5069/anti-PD1 n=12 mice. One-Way ANOVA with Tukey’s multiple comparisons test. g) Quantification of CD4^-^ as a percentage of CD3^+^ cells in the peripheral blood for the orthotopic NASH-HCC mice treated with AZD5069/anti-PD1 and IgG-control or anti-CD8α (left) and representative flow cytometry plots (right). IgG n=4 mice; anti-CD8α n=4 mice. Unpaired T-test. h) Quantification of CD3^+^ and CD4^+^ cells by IHC analysis in the tumours for the orthotopic NASH-HCC mice treated with AZD5069/anti-PD1 and IgG-control or anti-CD8α. IgG-control n=6 mice; anti-CD8α n=4 mice. i) Quantification of CD4^+^ as a percentage of CD3^+^ cells in tumours for the orthotopic NASH-HCC mice treated with AZD5069/anti-PD1 and IgG-control or anti-CD8α. IgG-control n=6 mice; anti-CD8α n=4 mice. j) Timeline schematic for the anti-CD8a depletion regime in the authothonous DEN/ALIOS model. k) Quantification of tumour burden (mm^3^) for DEN/ALIOS mice at day 284 for each treatment arm. Vehicle n=12 mice (data from Figure 3b); AZD5069/anti-PD1 + no isotype IgG control n=12 mice; AZD5069/anti-PD1 + isotype IgG control n=5 mice; AZD5069/anti-PD1 + anti-CD8a n=7 mice. One-way ANOVA with Tukey’s multiple comparisons test. l) Quantification of CD8^+^ counts/field in non-tumour and tumour for DEN/ALIOS mice from each treatment arm. AZD5069/anti-PD1 + isotype IgG control n=5 mice; AZD5069/anti-PD1 + anti-CD8a n=7 mice. Two-Way ANOVA with Tukey’s multiple comparisons test. m) Quantification of CD4^+^ counts/field in non-tumour and tumour for DEN/ALIOS mice from each treatment arm. AZD5069/anti-PD1 + isotype IgG control n=5 mice; AZD5069/anti-PD1 + anti-CD8a n=7 mice. Two-Way ANOVA with Tukey’s multiple comparisons test.

**Supplementary Data Figure 5.**
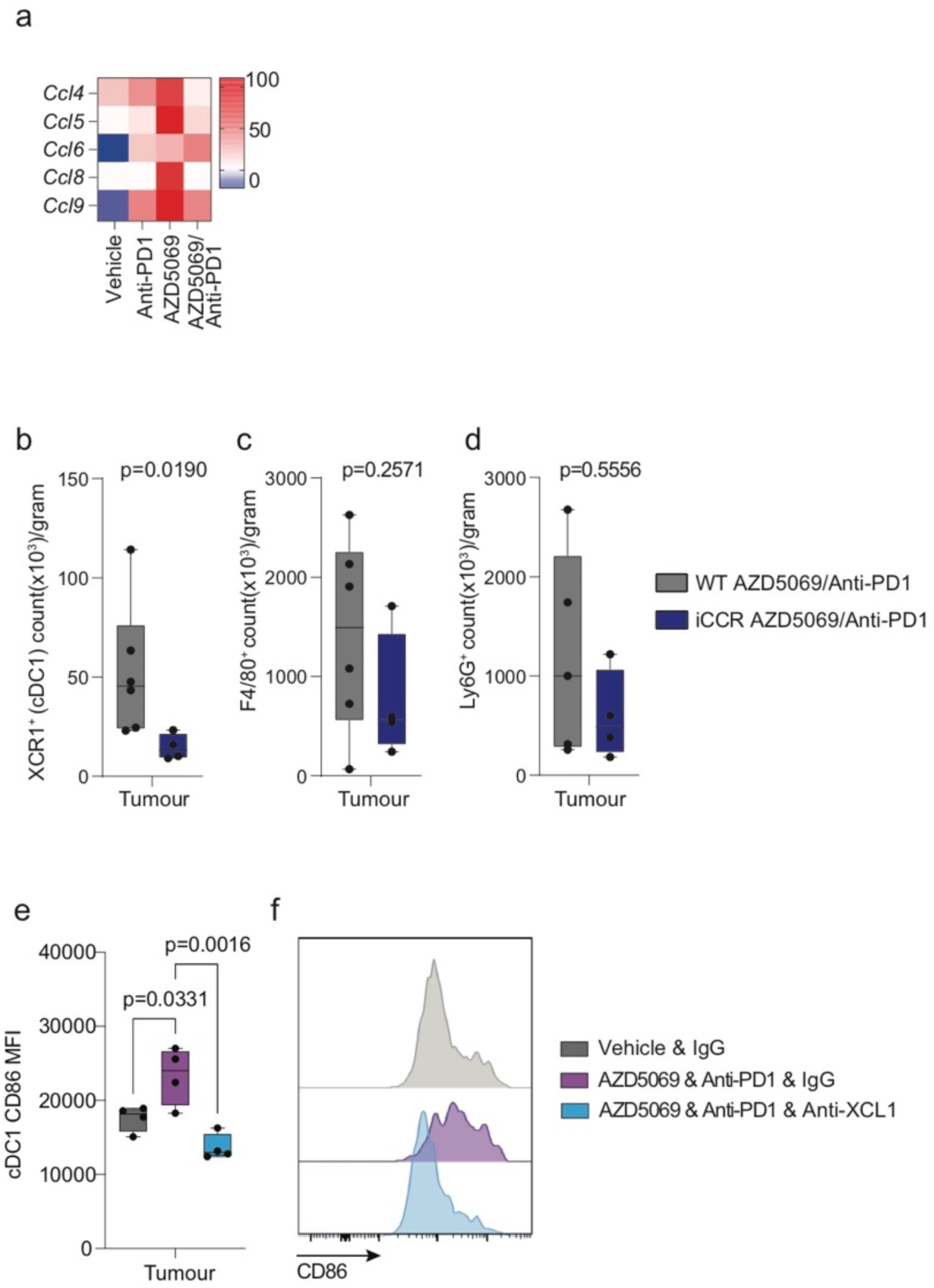
Dendritic cells contribute to combined AZD5069 & anti-PD1 anti-tumour phenotype. a) Heatmap showing row-scaled expression of *Ccl* chemokine transcripts, related dendritic cell recruitment, in DEN/ALIOS treated groups. Data are from bulk tumour RNA-Seq. b) Flow cytometric quantification of XCR1^+^ cDC1 counts (x10^3^)/gram in tumours of WT or iCCR genotype DEN/ALIOS mice treated with AZD5069/anti-PD1 at day 284. WT n=6. iCCR n=4. Mann-Whitney *U*-test. c) Flow cytometric quantification of F4/80^+^ macrophage counts (x10^3^)/gram in tumours of WT or iCCR genotype DEN/ALIOS mice treated with AZD5069/anti-PD1 at day 284. WT n=6. iCCR n=4. Mann-Whitney *U*-test. d) Flow cytometric quantification of Ly6G^+^ neutrophil counts (x10^3^)/gram in tumours of WT or iCCR genotype DEN/ALIOS mice treated with AZD5069/anti-PD1 at day 284. WT n=6. iCCR n=4. Mann-Whitney *U*-test. e) Flow cytometric quantification CD86 median fluorescence intensity (MFI) of XCR1^+^ cDC1s from tumuour of orthtopic NASH-HCC mice. Vehicle n=4. AZD5069/anti-PD1 and IgG n=4. AZD5069/anti-PD1 and Anti-XCL1 n=4. One-Way ANOVA with Tukey multiple comparisons test. f) Representative histogram plot of CD86 median fluorescence intensity (MFI) of XCR1^+^ cDC1s from tumuour of orthtopic NASH-HCC mice. Vehicle n=4. AZD5069/anti-PD1 and IgG n=4. AZD5069/anti-PD1 and Anti-XCL1 n=4. Dots in **Supplementary Data Fig. 6b-e**, represent individual mice.

**Supplementary Data Figure 6.**
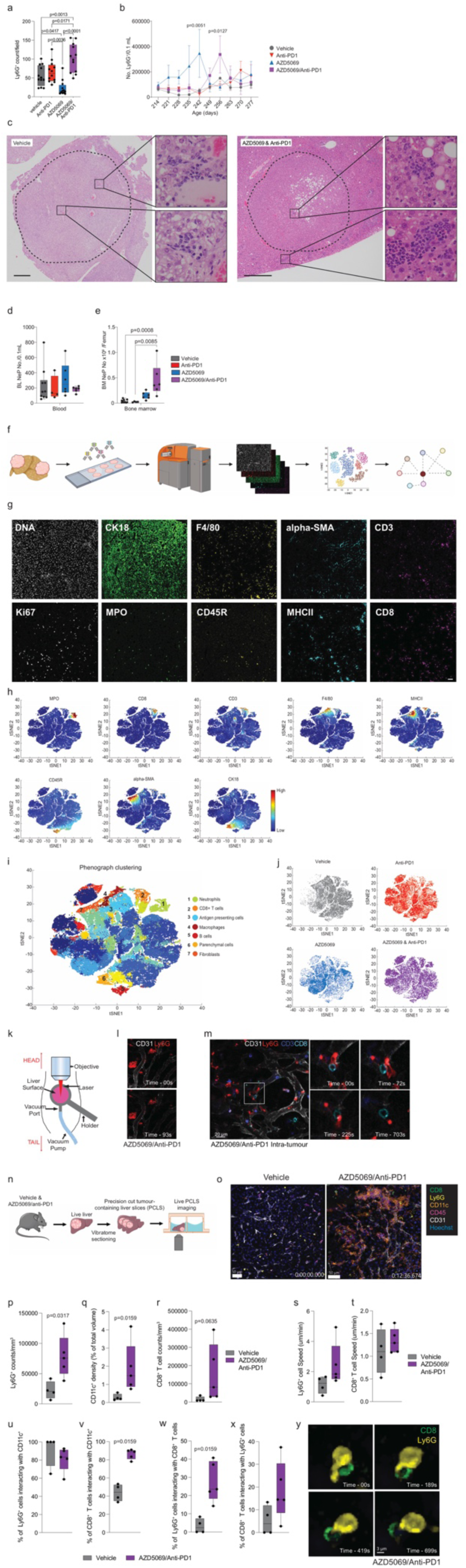
Imaging mass cytometry and live cell imaging reveals intra-tumour proliferating neutrophils associate and interact with CD8^+^ T cells and antigen presenting cells. a) Quantification of Ly6G^+^ counts/field in livers for the orthotopic NASH-HCC mice for each treatment arm. Vehicle n=12 mice; anti-PD1 n=12 mice; AZD5069 n=12 mice; AZD5069/anti-PD1 n=12 mice. One-Way ANOVA with Tukey’s multiple comparisons test. b) Flow cytometric quantification of the number of Ly6G^+^/0.1 mL peripheral blood from serial tail bleed analysis for DEN/ALIOS mice between day 214 and day 277. Vehicle: n=6 mice at all time-points except day 270 where n=5 mice. Anti-PD1: day 214-263 n=5 mice; day 270 n=3 mice; day 277 n=4 mice. AZD5069: n=5 mice at all time-points except day 242 where n=4 mice. AZD5069/anti-PD1: n=5 mice at all time-points except day 277 where n=4 mice. Two-way ANOVA with Tukey’s multiple comparisons test. Error bars represent Mean ± SEM. c) Representative images of H&E staining in livers for DEN/ALIOS Vehicle and AZD5069/anti-PD1 treated mice. Clusters of neutrophils with a banded morphology are enlarged in AZD5069/anti-PD1 treated mice. Scale bar = 1,000 µm. n=9 mice. d) Flow cytometric quantification of NeP count in 0.1ml blood for DEN/ALIOS mice for each treatment arm at day 284. Vehicle n=6 mice; anti-PD1 n=4 mice; AZD5069 n=6 mice; AZD5069/anti-PD1 n=6 mice. Two-way ANOVA with Sidak’s multiple comparisons test. e) Flow cytometric quantification of NeP count per femur for DEN/ALIOS mice for each treatment arm at day 284. Vehicle n=6 mice; anti-PD1 n=4 mice; AZD5069 n=6 mice; AZD5069/anti-PD1 n=6 mice. Two-way ANOVA with Sidak’s multiple comparisons test. f) Schematic of Fluidigm Hyperion imaging mass cytometry pipeline, including preparation of tumour tissue microarray, metal conjugated antibody staining, imaging mass cytometry and downstream analysis including tSNE and neighbourhood analysis performed using histoCAT. g) Representative images of hyperion staining for DNA intecalator, CK18, F4/80, alpha-SMA, CD3, Ki67, MPO, CD45R, MHCII and CD8. Scale bar = 100 µm. n=6 tumours. h) tSNE plots showing clustering of MPO, CD8, CD3, F4/80, MHCII, CD45R, alpha-SMA and CK18 positive cells. i) tSNE plot showing phenograph clustering of cell populations across all images and treatment groups. j) tSNE plots showing the contribution of individual treatment groups to clustering k) Schematic for DEN/ALIOS model liver intravital microscopy (IVM) set-up. l) Representative images for intra-tumour extravascular Ly6G^+^ clusters by IVM for DEN/ALIOS mice treated with AZD5069/anti-PD1 at day 284. m) Representative images for intra-tumour IVM for DEN/ALIOS mice treated with AZD5069/anti-PD1 at day 284 (right). Data are representative of n=1 mouse. Scale bar = 20 µm. n) Schematic for DEN/ALIOS model precision cut tumour-containing liver slice (PCLS) microscopy set-up. o) Representative images of live cell imaging of CD8, Ly6G, CD11c, CD45, CD31 and Hoechst in PCLS generated from DEN/ALIOS mice treated with either vehicle or AZD5069/anti-PD1 at day 284. Scale bar = 50 µm. p) Quantification of Ly6G^+^ counts/mm^3^ in PCLS generated from DEN/ALIOS mice treated with either vehicle or AZD5069/anti-PD1 at day 284. Vehicle n=4 mice. AZD5069/anti-PD1 n=5 mice. Mann-Whitney *U*-test. q) Quantification of CD11c^+^ density as a percentage of total volume in PCLS generated from DEN/ALIOS mice treated with either vehicle or AZD5069/anti-PD1 at day 284. Vehicle n=4 mice. AZD5069/anti-PD1 n=5 mice. Mann-Whitney *U*-test. r) Quantification of CD8^+^ T cell counts/mm3 in PCLS generated from DEN/ALIOS mice treated with either vehicle or AZD5069/anti-PD1 at day 284. Vehicle n=4 mice. AZD5069/anti-PD1 n=5 mice. Mann-Whitney *U*-test. s) Quantification of Ly6G^+^ speed in µm/min in PCLS generated from DEN/ALIOS mice treated with either vehicle or AZD5069/anti-PD1 at day 284. Vehicle n=4 mice. AZD5069/anti-PD1 n=5 mice. t) Quantification of CD8^+^ T cell speed in µm/min in PCLS generated from DEN/ALIOS mice treated with either vehicle or AZD5069/anti-PD1 at day 284. Vehicle n=4 mice. AZD5069/anti-PD1 n=5 mice. u) Quantification of the percentage of Ly6G^+^ cells interacting with CD11c^+^ cell surface in PCLS generated from DEN/ALIOS mice treated with either vehicle or AZD5069/anti-PD1 at day 284. Vehicle n=4 mice. AZD5069/anti-PD1 n=5 mice. v) Quantification of the percentage of CD8^+^ T cells interacting with CD11c^+^ cell surface in PCLS generated from DEN/ALIOS mice treated with either vehicle or AZD5069/anti-PD1 at day 284. Vehicle n=4 mice. AZD5069/anti-PD1 n=5 mice. Mann-Whitney *U*-test. w) Quantification of the percentage of Ly6G^+^ cells interacting with CD8^+^ T cells in PCLS generated from DEN/ALIOS mice treated with either vehicle or AZD5069/anti-PD1 at day 284. Vehicle n=4 mice. AZD5069/anti-PD1 n=5 mice. Mann-Whitney *U*-test. x) Quantification of the percentage of CD8^+^ T cells interacting with Ly6G^+^ cells in PCLS generated from DEN/ALIOS mice treated with either vehicle or AZD5069/anti-PD1 at day 284. Vehicle n=4 mice. AZD5069/anti-PD1 n=5 mice. y) Representative images of a Ly6G^+^ neutrophil and CD8^+^ T cell interacting in PCLS generated from DEN/ALIOS mice treated with AZD5069/anti-PD1 at day 284. Scale bar = 3 µm. Dots in **Supplementary Data Fig. 6a, d, e, p-x** represent individual mice.

**Supplementary Data Figure 7.**
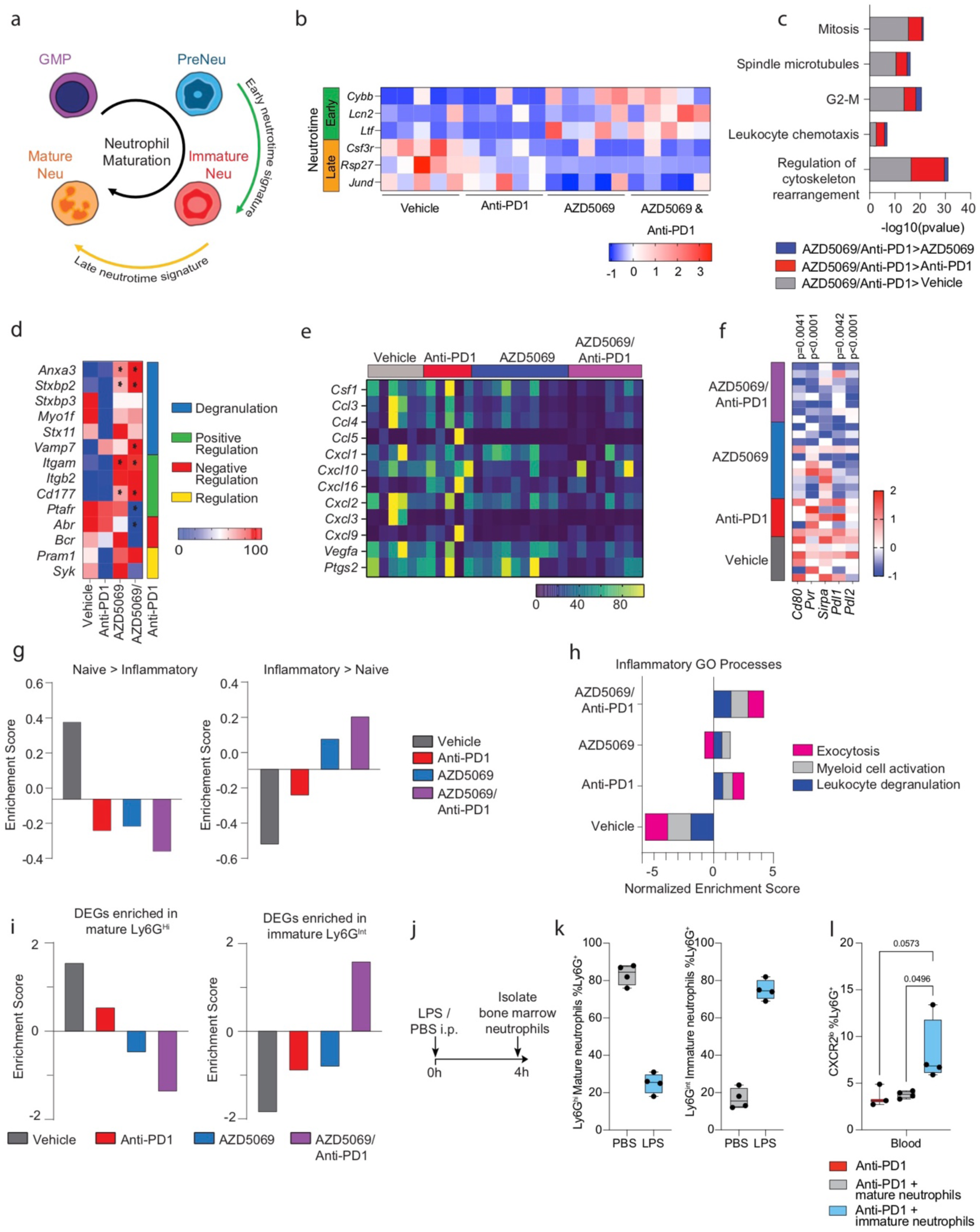
Combined AZD5069 & anti-PD1 therapy promotes an immature inflammatory tumour-associated neutrophil. a) Schematic depicting neutrophil maturation stages classified according to early and late neutrotime signatures (as described by Grieshaber-Bouyer *et al.*). b) Heatmap showing normalised expression of selected early and late neutrotime genes in neutrophils isolated from treated orthtoptic NASH-HCC mice at day 28. Data are from RT-PCR of isolated tumour neutrophils. N=5 neutrophils isolations per group. c) GSEA for process networks enriched in DEN/ALIOS TANs from AZD5069/anti-PD1 treated compared with Vehicle, anti-PD1 and AZD5069. Data are from bulk Ly6G^+^ RNA-Seq. d) Heatmap showing row-scaled expression of genes associated with neutrophil degranulation GO Processes (0043312, 0043313, 0043314, 0043315) for DEN/ALIOS mice TANs. Data are from bulk Ly6G^+^ TANs analysed by RNA-Seq. e) Heatmap showing row-scaled expression of genes associated with a pro-tumour neutrophil phenotype for DEN/ALIOS TANs from each treatment arm. Data are from bulk Ly6G^+^ TANs analysed by RNA-Seq. f) Heatmap showing row-scaled expression of immune checkpoint genes for DEN/ALIOS TANs. Data are from bulk Ly6G^+^ TANs analysed by RNA-Seq. g) GSEA from DEGs increased in neutrophils isolated from PBS-control (left) and LPS-treated mice (right) for TANs isolated from DEN/ALIOS mice for each treatment arm. Data are from bulk Ly6G^+^ TANs analysed by RNA-Seq. h) GSEA for inflammatory GO Processes enriched in LPS-treated compared with PBS-control mice for TANs isolated from DEN/ALIOS mice for each treatment arm. Data are from bulk Ly6G^+^ TANs analysed by RNA-Seq. i) GSEA for DEGs upregulated in LPS-treated peripheral blood, compared with PBS-controls, by mature Ly6G^Hi^ neutrophils (left) and immature Ly6G^int^ neutrophils (right) for DEN/ALIOS TANs. DEGs identified from bulk hepatic Ly6G^+^ neutrophils from PBS-control and LPS-treated mice (Mackey *et al.,* 2021) analysed by RNA-Seq, and GSEA from from bulk Ly6G^+^ TANs analysed by RNA-Seq. j) Schematic showing timeline of treatment of WT mice with LPS or PBS for harvesting of immature and mature neutrophils. k) Flow cytometric quantification of Ly6G^hi^ and Ly6G^int^ neutrophils as a percentage of total Ly6G^+^ bone marrow cells from LPS and PBS treated mice. LPS n=4. PBS n=4. l) Flow cytometric quantification of CXCR2^Lo^ neutrophils as a percentage of Ly6G^+^ neutrophils from the blood of orthotopic NASH-HCC, neutrophil/anti-PD1 therapy mice at day 28. Anti-PD1 n=3. Anti-PD1/Mature neutrophils n=4. Anti-PD1/Immature neutrophils n=4. Bulk DEN/ALIOS Ly6G^+^ TAN RNA-Seq data in **Supplementary Data Fig. 7b-h**: Vehicle n=6 mice; anti-PD1 n=5 mice, AZD5069 n=10 mice; AZD5069 n=8 mice. Bulk hepatic Ly6G^+^ neutrophil RNA-Seq data in **Supplementary Data Fig. 7** from Mackey *et al.,* 2021. Dots in **Supplementary Data Fig. 7j-q** represent individual mice.

**Supplementary Movie – Neutrophils associate and interact with CD8^+^ T cells and CD11c^+^ cells.**

Representative time-lapse movies of tumour bearing precision cut liver slices generated from livers from DEN/ALIOS mice treated with either vehicle or AZD5069/anti-PD1 at day 284. Samples were stained for CD8 (green), Ly6G (yellow), CD11c (orange), CD45 (purple), CD31 (white) and Hoechst (blue) and imaged for 15-40min in a full incubation chamber at 37°C with 5% CO2. Scale bar = 50 µm.

**Supplementary table 1:**
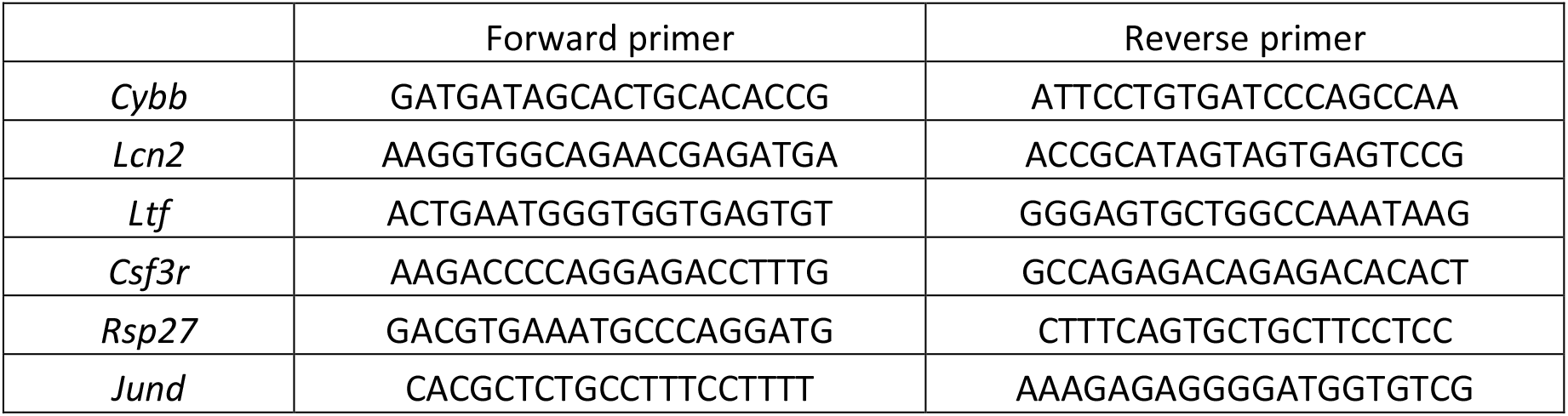
List of mouse primer sequences used for qRT-PCR analysis.

